# Clonotype specificity distinguishes epitope-tailored and permissive somatic hypermutation

**DOI:** 10.64898/2026.10.10.758199

**Authors:** Vishal N. Rao, Bruno Bonnettaz, Alesandro Civljak, Anass Abbad, Florian Krammer, Camila H. Coelho, Goran Bajic

## Abstract

Identifying antibody features that reliably encode epitope specificity poses a major barrier towards modeling antigen-antibody interactions. Affinity maturation offers a unique opportunity to probe the molecular basis of antigen-antibody interactions by revealing how individual somatic hypermutations (SHM) refine antibody binding. Using SARS-CoV-2 spike-specific germinal center B-cell repertoires, we show that antibodies harboring multiple SHMs that are restricted to their respective clonotypes (clonotype-restricted SHMs) preferentially recognize cryptic epitopes. We structurally characterized three such antibodies: two recognized a cryptic epitope on the receptor-binding domain whereas a third bound an N-terminal domain epitope that is almost entirely occluded in prefusion spike. In contrast, SHMs shared across multiple clonotypes using the same V gene recur across antibodies targeting diverse antigens but generally fail to enhance affinity when introduced into diverse antibody backgrounds. Together, these findings define a spectrum of SHM behavior, distinguishing epitope-tailored mutations from broadly recurrent but largely permissive substitutions.

**One Sentence Summary:** The extent to which a somatic hypermutation is shared across antibody clonotypes distinguishes epitope-tailored mutations associated with cryptic epitope recognition, from recurrent, largely permissive substitutions.

## INTRODUCTION

Antigen-antibody interactions constitute a dynamic system of host-pathogen co-evolution at the molecular level. During affinity maturation, somatic hypermutation (SHM) generates antibody variants that undergo selection for improved antigen recognition^1,2^, while pathogens, particularly viruses, accumulate sequence changes that can reduce antibody binding and enable immune escape^3,4^. This reciprocal process reflects opposing selective pressures: antibody diversification and selection optimize recognition within a given antigenic context, whereas viral evolution favors variants that evade existing humoral immunity. Understanding this molecular arms race requires defining the principles that govern how antibody sequence variation translates into antigen recognition.

One strategy to uncover these principles is to examine how antibodies gain affinity during germinal center-mediated maturation. SHM introduces mutations into antibody variable genes, while selection in the context of the cognate antigen enriches variants with improved binding^5^. Through iterative rounds of diversification and selection, germinal centers explore mutational solutions that enhance antigen recognition^6,7^. Comparing antibody lineages at different stages of this process can therefore reveal the structural and biochemical constraints that shape antigen–antibody interactions. This raises a central question: can patterns of SHM acquired during affinity maturation predict the nature of antigen–antibody interactions and, consequently, the characteristics of the targeted epitope?

Associations between antibody sequence features and epitope specificity have traditionally been examined at the level of germline gene usage^8–11^. Although germline biases can reveal broad preferences for particular epitopes, they provide limited resolution into the specific molecular determinants of antigen–antibody interactions, including the residues involved and the physicochemical nature of their interactions. SHMs selected at individual positions may provide more granular information. In addition to their location within the antibody, the chemical nature of the resulting amino acid substitutions can reveal how affinity maturation adapts the paratope to the local biochemical environment of an epitope. For example, we previously showed that rare clonotype-restricted mutations in the severe acute respiratory syndrome coronavirus 2 (SARS-CoV-2)-specific public antibody M15 involved recurrent polar-to-hydrophobic substitutions that facilitated recognition of a hydrophobic cleft within a cryptic S2 epitope^12^. Similar examples of epitope-specific adaptation through SHM have been described in other antibody responses^9,13,14^, suggesting that selected mutations can encode structural solutions tailored to specific epitope features.

However, the mere presence of an SHM does not imply that it directly enhances affinity. GC B cells acquire SHM through activation-induced cytidine deaminase (AID) during proliferation in the dark zone, whereas affinity-based selection occurs subsequently in the light zone^2,6,15,16^. Selection therefore acts on B cell receptor (BCR) variants carrying combinations of mutations rather than evaluating individual SHMs in isolation. As a result, mutations that are neutral or even mildly unfavorable can persist within positively selected clones, and a substantial fraction of acquired SHMs do not measurably enhance affinity^17^. This mutational noise complicates efforts to infer epitope characteristics from individual SHMs based on sequence information alone.

Previous efforts to model SHM acquisition, including the Synonymous 5-mer Functional (S5F) model and the phylogenetic codon-substitution model derived from GY94, have largely focused on the sequence determinants and evolutionary patterns of SHM^18–22^. However, understanding why particular mutations are selected within individual antibody lineages requires consideration of the antigenic context in which they arise. We previously showed that the frequency of a given SHM within a public antibody clonotype, relative to its frequency among other antibodies using the same V gene, can provide information about both its contribution to affinity maturation and the biochemical nature of the underlying antigen–antibody interactions^12^. We term this property clonotype specificity. Because clonotype specificity is derived from the observed distribution of mutations across antibody repertoires, it integrates the effects of both intrinsic SHM targeting and subsequent selection. Two key questions, however, remained unresolved: (i) can clonotype specificity predict recognition of cryptic epitopes beyond public antibody clonotypes, and (ii) conversely, what role do clonotype-shared SHMs play in antigen–antibody interactions, and do they consistently contribute to affinity enhancement?

In this study, we address these questions using spike-specific germinal center BCR clonotypes elicited following SARS-CoV-2 mRNA vaccination together with antigen–antibody structural data curated in Structural Antibody Database (SAbDab)^23^. We show that SARS-CoV-2 spike-specific antibody lineages that accumulate multiple clonotype-restricted SHMs preferentially recognize cryptic epitopes on the SARS-CoV-2 spike protein. In contrast, clonotype-shared SHMs recur not only among spike-specific antibodies but also across antibodies targeting unrelated antigens that use the same V gene, revealing a broader, “pan-antigenic” nature. These mutations frequently adopt conserved structural roles, including antigen and cognate-chain contacts, yet generally fail to enhance affinity when introduced into diverse antibody backgrounds. Collectively, these findings define distinct patterns of SHM behavior during affinity maturation. Frequently sampled, pan-antigenic SHMs appear to represent largely permissive mutational outcomes with rare affinity benefits, whereas rarer clonotype-restricted SHMs encode antigen- or epitope-specific solutions, including those required for recognition of cryptic epitopes. This framework links patterns of SHM across antibody repertoires to the structural constraints that shape antigen recognition.

## RESULTS

### Clonally expanded GC B cells induced by SARS-CoV-2 spike mRNA vaccination exhibit varying degrees of clonotype-specificity in SHM

To evaluate clonotype-specificity across distinct germinal center (GC) B cell lineages, we analyzed a dataset of SARS-CoV-2 spike–specific GC BCR sequences obtained following ancestral mRNA vaccination (**Figure 1A**). Leveraging clonotypes pre-assigned by Kim and colleagues^1^, we first restricted our analysis to clonal lineages comprising at least 20 clones to ensure sufficient sampling depth (n = 423 clonotypes). To focus on mutations propagated within sublineages, we defined frequent SHMs as those present in at least 30% of clones within a lineage, a threshold chosen to capture mutations that are effectively fixed within expanding sublineages. To assess clonotype-specificity within V genes, we further restricted our analysis to V genes that (i) harbored at least 10 frequent SHMs overall and (ii) were represented by at least 10 distinct clonotypes. For each qualifying V gene, clonotype-specificity of a given frequent SHM was defined as the number of clonotypes in the dataset sharing that mutation (**Figure 1A**). Across V genes, SHMs displayed a broad spectrum of clonotype specificities in both the complementarity-determining regions (CDRs) and framework regions (FWRs) (**Figure 1B, C**).

**Figure 1:**
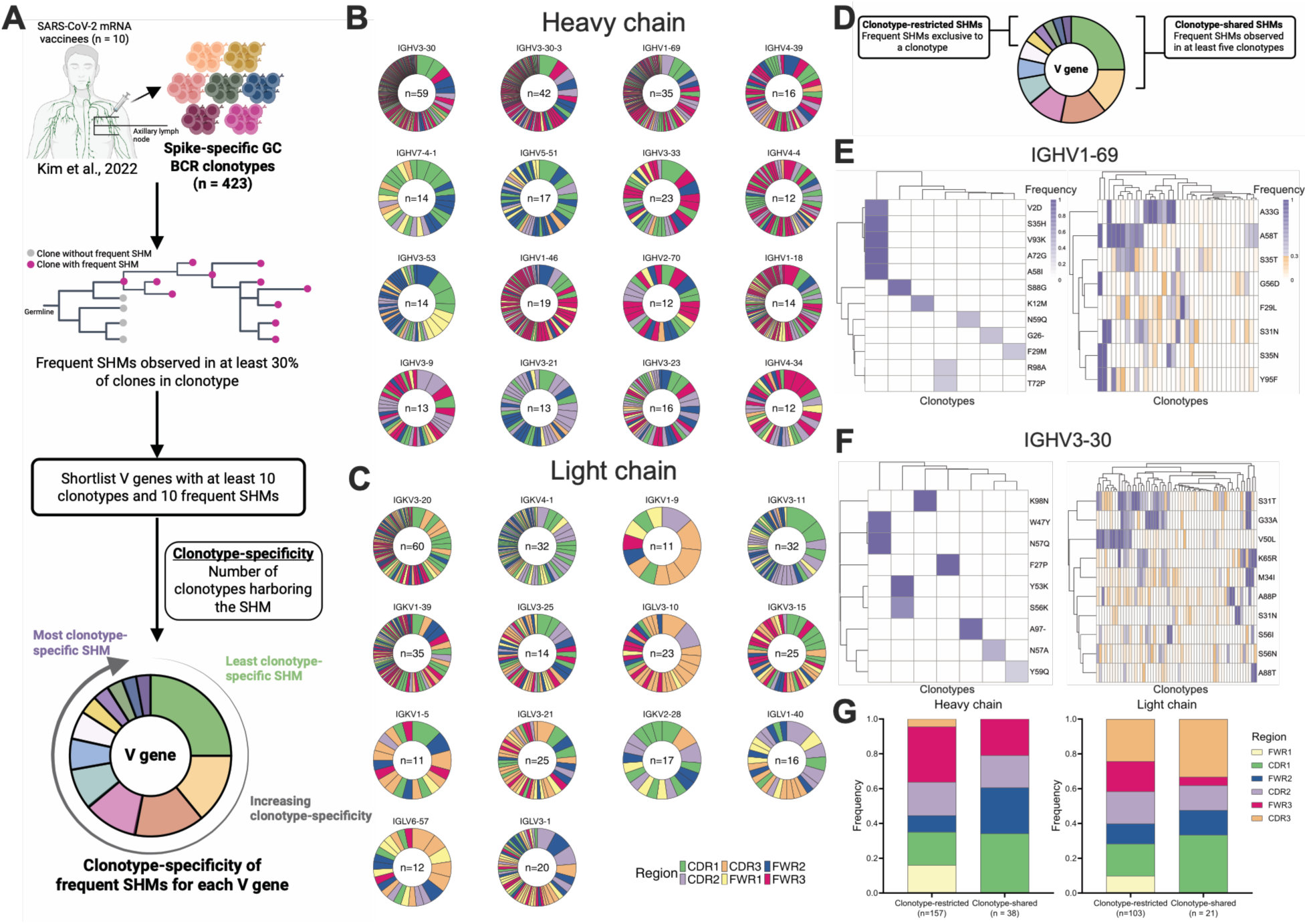
Spike-specific GC BCR clonotypes display SHMs with a wide range of clonotype-specificities. (**A**) Schematic representation of the method used to compute clonotype specificity of frequent SHMs for each V gene in the spike-specific GC BCR repertoire retrieved from Kim and colleagues^1^. The color of the tips in the representative clonal tree indicates the presence of the given SHM in the clone. Each segment in the representative donut plot is an SHM arranged in increasing order of clonotype specificity clockwise. (**B-C**) Donut plots representing the clonotype specificity for each frequent SHM identified for each selected (**B**) heavy and (**C**) light chain V gene. Each segment represents an SHM, with the size of the segment proportional to the number of clonotypes harboring the SHM. The mutations are arranged from most clonotype-sharing to most clonotype-specific clockwise. The color of each segment indicates the position of the mutation (CDR/FWR). (**D**) Schematic representing the method for shortlisting clonotype-restricted and clonotype-shared SHMs. (**E-F**) Heatmaps depicting the frequency of clonotype-restricted (left) and clonotype-shared SHMs (right) in (**E**) IGHV1-69 and (**F**) IGHV3-30 spike-specific BCR clonotypes. The rows represent clonotype-restricted/clonotype-shared SHMs and the columns represent clonotypes. (**G**) Frequency of clonotype-restricted and clonotype-shared SHMs in different CDR/FWRs in heavy (left) and light chain (right) clonotypes, compiled across all shortlisted V genes. The CDR/FWR region is represented by the color of each segment.

We then focused on SHMs at the extremes of the clonotype-specificity spectrum; clonotype-shared SHMs were defined as frequent mutations observed in at least five clonotypes using the same V gene, whereas clonotype-restricted SHMs were those exclusive to a single clonotype (**Figure 1D-F**). Thus, clonotype-shared SHMs likely represent mutations with high likelihood of being targeted by AID, but low epitope-specific selection. Clonotype-restricted SHMs, on the other hand, likely represent those with low to moderate targeting, but high epitope-specific selection. This enables us to assess factors influencing selection of a mutation, largely independent of AID targeting biases. We compared the distribution of clonotype-shared and clonotype-restricted SHMs across FWR1, CDR1, FWR2, CDR2, FWR3, and the initial residues of CDR3 to determine whether these mutations were differentially distributed across CDRs and FWRs, given their distinct roles in antigen interactions. Notably, in both heavy and light chains, clonotype-restricted SHMs were enriched in FWR3, whereas clonotype-shared SHMs were enriched in CDR2 (**Figure 1G**).

### Antibodies with clonotype-restricted SHM target cryptic epitopes on SARS-CoV-2 spike

We observed multiple clonotypes that accumulated several clonotype-restricted SHMs (**Figure 1D**), prompting us to investigate whether these clonotypes preferentially recognize cryptic epitopes on the SARS-CoV-2 spike protein. We previously identified a public antibody clonotype that uses clonotype-restricted SHMs to recognize a cryptic spike epitope, motivating this hypothesis^12^. Kim and colleagues^1^ previously assessed spike binding by selecting one representative clone from each clonotype identified by flow cytometry-based sorting and testing its binding to the ancestral SARS-CoV-2 spike by Enzyme-linked immunosorbent assay (ELISA). They classified a clonotype as spike-positive when the representative clone showed detectable spike binding. Clonotype-restricted SHMs occurred more frequently in clonotypes classified as spike-negative by ELISA (**Figure S1**), suggesting that some of these lineages may bind spike below the detection threshold of the assay or recognize non-spike antigens.

To determine whether accumulation of multiple clonotype-restricted SHMs is indicative of cryptic epitope recognition among spike-positive clonotypes, we identified clonotypes harboring at least three clonotype-restricted SHMs (**Figure 2A**). For each clonotype, we selected the most mutated clone containing all identified clonotype-restricted SHMs and expressed it as a fragment antigen-binding (Fab) version of the monoclonal antibody (mAb). In two clonotypes, clonotype-restricted SHMs were distributed across two distinct sublineages (**Figure S2A, B**); in these cases, the most mutated clone from each relevant sublineage was selected for recombinant expression and purification (CR15-16; CR17-18).

**Figure 2:**
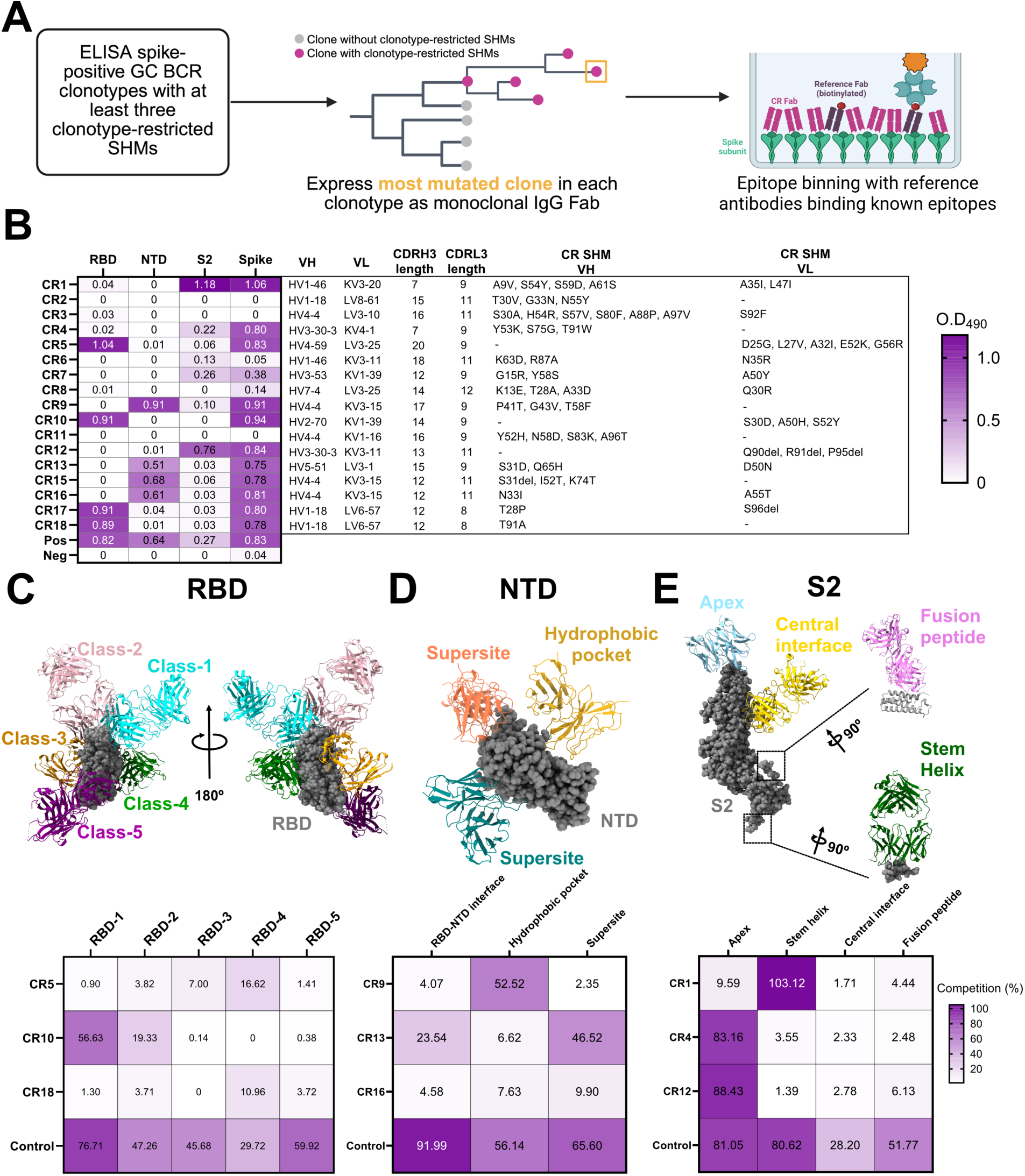
Spike subunit specificity and epitope binning of CR Fabs. (**A**) Schematic workflow for selection of clones for Fab expression and subsequent epitope binning, based on the number of clonotype-restricted SHMs identified per clonotype. For epitope binning, biotinylated reference Fabs targeting defined epitopes were incubated with CR Fabs added at a 20-fold molar excess. (**B**) ELISA analysis of CR Fab binding to full-length spike protein to determine subunit specificity. The table on the right summarizes sequence features of the corresponding clonotypes, including heavy and light chain V gene usage, CDR3 length, and the clonotype-restricted SHMs identified in each clonotype. (**C–E**) Epitope binning assays between CR Fabs and reference Fabs recognizing defined epitopic regions within (**C**) RBD, (**D**) NTD, and (**E**) S2. The epitopic regions targeted by reference Fabs are illustrated above. Based on the subunit specificities assigned in (**B**), CR Fabs were competed against biotinylated reference Fabs, with CR Fabs present at a 20-fold molar excess. Competition was quantified as the percentage decrease in biotinylation signal of the reference Fab upon addition of the CR Fab.

We generated a panel of 18 Fabs, hereafter termed ‘clonotype-restricted (CR) Fabs’ (CR1–CR18) and assessed their spike subunit specificity by ELISA. We subsequently excluded CR14 after clonal lineage inference resolved an ambiguous pair of polymorphisms that we had initially classified as SHMs. Among the remaining Fabs, four (CR5, CR10, CR17, CR18) bound the receptor-binding domain (RBD), four (CR9, CR13, CR15, CR16) bound the N-terminal domain (NTD), and three (CR1, CR4, CR12) bound the S2 subunit of spike (**Figure 2B**).

Despite originating from clonotypes previously classified as spike-positive, three Fabs exhibited only weak binding (CR6, CR7, CR8), whereas three others (CR2, CR3, CR11) showed no detectable binding by ELISA (**Figure 2B**). To investigate these discrepancies, we recombinantly expressed CR2, CR3 and CR11 as full-length IgG1s together with the corresponding antibodies from the same clonotypes that Kim and colleagues had previously reported as spike-positive. In all three clonotypes, the CR Fabs selected for this study had accumulated more SHMs relative to germline than the antibodies characterized by Kim and colleagues (**Figure S3A, B, C**). Neither antibody from the CR2 and CR11 clonotypes bound spike, whereas the antibody characterized by Kim and colleagues from the CR3 clonotype retained detectable binding by ELISA (**Figure S3D, F, H**). We further confirmed these results by biolayer interferometry: the CR3 counterpart characterized by Kim and colleagues was the only mAb with detectable spike binding, although the interaction was of low affinity (**Figure S3E, G, I**).

Comparison of the two CR3 antibodies revealed both shared and clone-specific SHMs. CR3 uniquely contained Q3K and V72M in the heavy chain and G94N and V98I in the light chain, whereas the antibody characterized by Kim and colleagues uniquely contained S28F, V72I, Q78H in the heavy chain and H96Q in the light chain (**Figure S3J**). Notably, both antibodies contained all clonotype-restricted SHMs associated with the CR3 clonotype, indicating that these mutations alone cannot account for the loss of spike binding by CR3.

To determine whether the spike-binding CR Fabs recognized cryptic epitopes, we performed epitope binning by competing each Fab against a panel of biotinylated reference Fabs targeting defined epitopic regions across the spike domains (**Figure 2C–E**). Because CR15/CR16 and CR17/CR18 originated from the same clonotypes, we selected one representative Fab from each pair (CR16 and CR18, respectively) to avoid redundancy. Among the RBD-binding Fabs, CR10 strongly competed with the RBD-1 reference antibody, whereas CR5 weakly competed with RBD-3 and RBD-4, and CR18 weakly competed with RBD-4. Among the NTD-binding Fabs, CR9 competed with antibodies recognizing the RBD–NTD interface, whereas CR13 competed with an antibody recognizing a hydrophobic NTD epitope. In contrast, CR16 showed minimal competition with all reference antibodies (**Figure 2D**). Among the S2 binders, CR1 strongly competed with a stem-helix–directed antibody, whereas CR4 and CR12 competed with apex-directed antibodies (Figure 2E).

Collectively, epitope binning identified three CR Fabs (CR5, CR16, CR18) that showed limited competition with the reference antibody panel. Because this panel encompasses many commonly targeted regions of spike, these results suggest that the three CR Fabs may recognize distinct epitopes that are poorly represented among the established antibody specificities, including potential cryptic epitopes.

### CR antibodies recognize cryptic epitopes on sarbecoviruses

To determine whether the CR antibodies with weak or no competition against reference antibodies recognize cryptic spike epitopes, we determined cryo-EM structures of the RBD-specific CR5 and CR18 and NTD-specific CR16 (**Table S1, Figure S4, S5, S6**). We structurally characterized CR5 and CR18 in complex with the RBD and non-competing Class-1 and Class-5 Fabs (**Figure 3A, C, S4, S5, S7A,B**). We resolved the CR5-RBD interface at sufficient resolution to define residue-level interactions. CR5 recognizes an epitope centered on the 377–386 strand-loop-helix region, with an additional hydrophobic patch comprising L390, F392, and L517 (**Figure 3A, S8A**). The CR5 light chain dominates the interface, engaging the RBD through all three CDRs and framework residue Y2, whereas HCDR3 forms a more restricted interaction area. Heavy-chain Y108 contacts C379, while D110 interacts with S383, which also contacts light-chain Q30 (**Figure S8A**). The CR5 epitope closely overlaps the cryptic epitope recognized by CR3022 and its affinity-matured variants^24^, including the conserved 377–386 loop and the secondary 429/515–517 region (**Figure S8A-C, S9A-E**). However, CR5 approaches the epitope at an angle approximately 45° different from that of CR3022 (**Figure S7A**).

**Figure 3:**
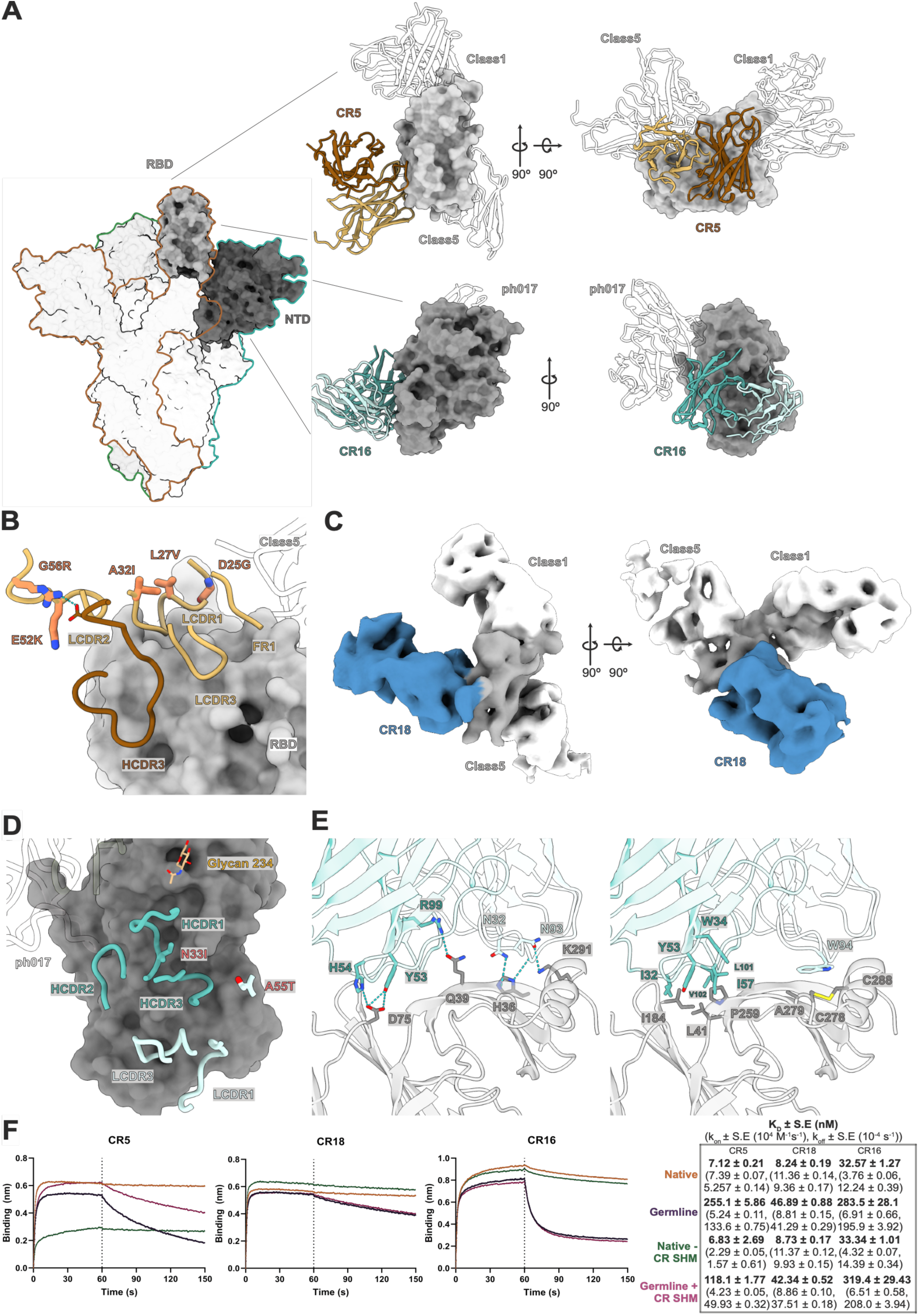
Cryo-EM structural analysis of CR Fabs targeting cryptic epitopes. **(A)** Surface representation of the trimeric spike protein with one RBD and NTD domains from adjacent protomers highlighted in grey. (top-right) Structure of the CR5 antibody (heavy chain in maroon, light chain in beige) bond to the buried face of the RBD. Class 1 and class 5 antibody models are transparent. (bottom-right) Structure of the CR16 antibody (heavy chain in blue-green, light chain in light blue) bound to the buried face of the NTD. Supersite antibody (ph017) model is transparent. **(B)** Interface between CR5 and its cryptic epitope in RBD, with clonotype-restricted SHMs in CR5 labelled and colored in orange. **(C)** Moderate resolution cryo-EM map of CR18 (blue) in complex with RBD (grey) bound by non-competing Class-1 and Class-5 Fabs (transparent). **(D)** Binding footprint of CR16 heavy and light chains interacting with a buried epitope in NTD. Clonotype-restricted SHMs in CR16 are highlighted and labelled in red. **(E)** Residue interactions between CR16 (blue-green and light blue) and the buried epitope on NTD (light grey). All interacting residues are highlighted and labelled. The disulphide bond between C278 and C288 of NTD that is contacted by W94 in LCDR3 is highlighted in yellow. **(F)** Binding kinetics of (left to right) CR5 and CR18 with RBD, and CR16 with NTD. Each color corresponds to an expressed variant of the antibody. The dotted line at 60s represents the end of association and start of dissociation. Computed dissociation constant (K_D_) values along with the association (k_on_) and dissociation rates (k_off_) for each antibody variant are shown in the table on the right.

All five clonotype-restricted SHMs in CR5 occur in the IGLV3-25 light chain (**Figure 2B, 3B**). D25G and L27V primarily alter local LCDR1 geometry and packing, potentially optimizing the presentation of the antigen-contacting residues in LCDR1. In contrast, A32I, E52K, and G56R introduce packing or electrostatic interactions that rigidify the broader paratope. A32I introduces a bulkier hydrophobic side chain at a Vernier-zone^25^ framework position underlying the CDR loops, allowing it to pack simultaneously against LCDR1 and LCDR2 to stabilize the relative positioning of antigen-contacting Y31 and D50. E52K may further constrain LCDR2 through an interaction with framework E49, whereas G56R establishes an inter-chain interaction with HCDR3 residue D114 (**Figure 3B, S8A**). Thus, rather than creating new antigen contacts, the clonotype-restricted SHMs in CR5 appear to promote paratope pre-organization, consistent with a role for light-chain maturation in enabling recognition of a sterically restricted epitope^12^.

CR18 exhibited a binding mode similar to that of R1-26 ^26^ approaching the cryptic RBD epitope with a comparable heavy- and light-chain orientation (**Figure 3C, S7B**). Notably, CR18 and R1-26 both use the IGLV6-57 light-chain gene. Together, the CR5 and CR18 structures define a highly conserved cryptic RBD epitope. Superposition onto prefusion spike structures indicates that antibody access to this site requires substantial RBD opening and rotation, which relieve steric clashes with neighboring RBD and S2 regions and are favored when at least two RBD protomers adopt the up conformation^27^ (**Figure S10A**).

CR16 recognizes a distinct, underappreciated cryptic epitope on the sarbecovirus NTD that is largely buried in available prefusion spike structures. Nevertheless, CR16 binds the full-length ectodomain, indicating that the epitope can become accessible on the intact spike, potentially through local conformational breathing of the protomers or on shed S1 subdomain (**Figure 2B, S10B**). The discontinuous epitope comprises four major clusters distributed across both antibody chains. LCDR1 contacts H36/S37 near the NTD–RBD junction, whereas LCDR3 engages the C-terminal α-helix through K291 and the C278–C288 disulfide bond (**Figure 3D, E**). HCDR2 engages D75 and P259, while HCDR1/HCDR3 form a hydrophobic interaction network within a pocket between adjacent β-strands (**Figure 3E**).

CR16 contains two clonotype-restricted SHMs, N33I in HCDR1 and A55T in the light chain. N33I replaces a polar residue with a branched hydrophobic side chain and may promote a local loop rearrangement toward the center of the immunoglobulin fold, accommodating the NTD glycan at position 221 (**Figure 3D**). A55T may introduce a polar interaction with HCDR3 residue R103 and stabilize the HCDR3 conformation (**Figure 3D**). The clonally related antibody CR15 contains three additional clonotype-restricted mutations in the heavy chain, S31del, I52T, and K74T, that may further remodel and rigidify the CDR loops. S31del shortens the HCDR1 loop, whereas I52T may stabilize HCDR2; K74T may contribute indirectly through framework packing. Compared with BLN8, an antibody recently shown by Zhou and colleagues^28^ to recognize an overlapping NTD-8 epitope, CR16 extends its footprint toward a more buried surface of the NTD and approaches the epitope from an orientation compatible with the relative positioning of the adjacent RBD (**Figure S11A, B**).

While clonotype-restricted SHMs appeared to reorganize the paratope to accommodate cryptic epitopes rather than directly contact antigen, whether they contributed to affinity enhancement remains unclear. To test this, we generated four variants of CR5, CR16 and CR18: (i) the native version, (ii) a germline-reverted version with all mutations reverted to germline, (iii) the native antibody with only the clonotype-restricted SHMs reverted to germline and (iv) the germline-reverted antibody containing only the clonotype-restricted mutations. We measured their binding affinities to the cognate antigens (RBD and NTD) using biolayer interferometry (**Figure 3F**). All three native antibodies bound with higher affinities than their germline-reverted variants with 36-, 6- and 9-fold difference in K_D_ for CR5, CR18 and CR16, respectively, demonstrating substantial affinity maturation. However, reverting the clonotype-restricted SHMs in the native antibodies did not significantly reduce affinity, and introducing these mutations into the germline-reverted backgrounds did not significantly increase affinity (**Figure 3F**).

For CR5, however, the clonotype-restricted SHMs specifically affected association kinetics. Reverting these mutations in the native antibody significantly decreased the association rate (k_on_) whereas introducing them into the germline-reverted variant significantly increased k_on_; neither manipulation measurably affected the dissociation rate (k_off_). Together with the structural data, these results suggest that the five clonotype-restricted SHMs primarily reorganize the paratope in a manner that facilitates productive engagement of the cryptic epitope during association rather than stabilizing the final bound complex.

We next tested whether CR5, CR16 and CR18 neutralized ancestral SARS-CoV-2 in a live-virus neutralization assay. Of the three antibodies, only CR18 neutralized SARS-CoV-2 (IC_50_ = 2.52 μg/mL) (**Figure S12**), consistent with previously described IGLV6-57-encoded antibodies of this class^26^. Together, these findings suggest that clonotype-restricted SHMs can facilitate recognition of cryptic epitopes by optimizing paratope architecture without necessarily increasing net equilibrium affinity.

### Clonotype-shared SHMs are prevalent and exhibit similar functional roles in diverse antigen-antibody interactions

The association between accumulation of clonotype-restricted SHMs and cryptic epitope recognition observed across clonotypes prompted us to address the complementary question of how clonotype-shared SHMs shape antigen–antibody interactions. While clonotype-restricted mutations appeared to encode epitope-tailored structural solutions, clonotype-shared SHMs recur across independent clonotypes that use the same V gene and may therefore reflect more general features of affinity maturation.

Because clonotype-shared SHMs represent identical substitutions independently acquired across distinct spike-specific clonotypes (**Figure 1C**), we hypothesized that these mutations might also recur among spike-specific antibodies beyond our SARS-CoV-2-specific germinal center BCR dataset. To test this, we queried the Structural Antibody Database (SAbDab) for SARS-CoV-2 spike-specific antibody structures in complex with spike or its subdomains. We focused on antibodies using IGHV1-69 or IGHV3-30, two of the most frequently used functional V genes in the human antibody repertoire (**Figure 4A**). Clonotype-shared SHMs identified in the GC dataset frequently recurred among spike-specific antibodies using these V genes (**Figure 1C, 4B, C**).

**Figure 4:**
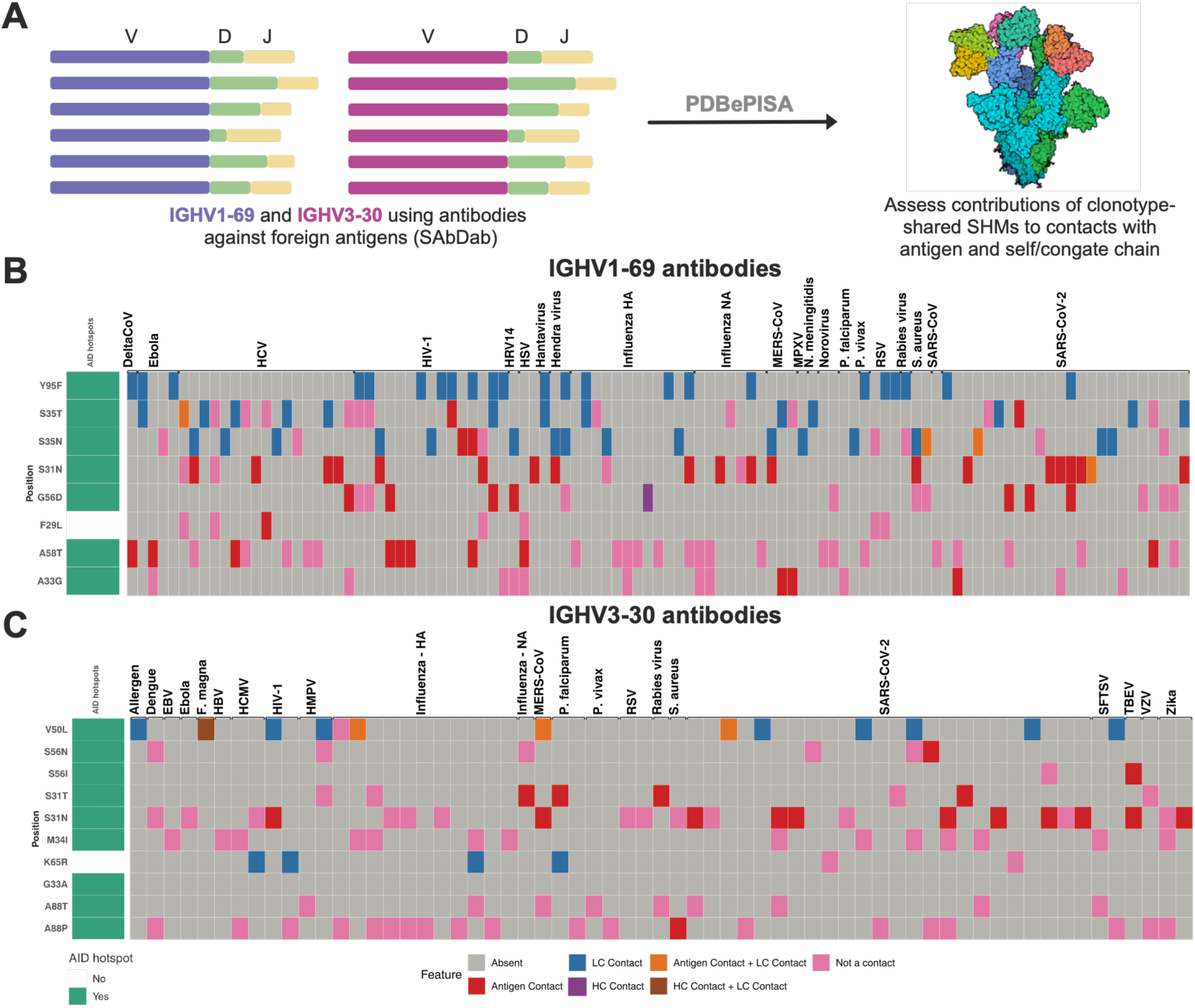
Functional role of clonotype-shared SHMs in antigen-antibody interactions. (**A**) Schematic representing the workflow employed to identify the role of clonotype-shared SHMs in antigen-antibody interactions, using the tool PDBePISA. (**B-C**) Heatmaps depicting the occurrence and function of clonotype-shared SHMs (rows) in making contacts with antigen, heavy chain and/or light chain, in the context of SARS-CoV-2 antibodies (center) and antibodies against other antigens, using the (**B**) IGHV1-69 or (**C**) IGHV3-30 genes. Each column represents an antigen-antibody structure retrieved from SAbDab. The heatmap on the left indicates whether the clonotype-shared SHMs are elicited at WRCY AID hotspot motifs.

The availability of structural data further enabled us to assess the functional contributions of these mutations. Using PDB-ePISA (Protein Data Bank Europe - Proteins, Interfaces, Structures and Assemblies) to map protein–protein contacts, we determined whether clonotype-shared SHMs formed interactions with antigen, heavy-chain, or light-chain residues. Although individual mutations participated in different types of interactions, each SHM exhibited a strikingly consistent functional role across independent antibodies (**Figure 4B, C**). Among IGHV1-69 spike-specific antibodies, S31N contributed to antigen contacts in all observed instances (7/7), Y95F consistently mediated light chain contacts, albeit at a low frequency (2/2), and A58T generally did not participate in direct contacts (4/5) (**Figure 4B**). Similarly, among IGHV3-30 antibodies, S31N frequently formed antigen contacts (7/10), V50L consistently mediated light chain contacts (5/5), whereas M34I (5/5), K65R (2/2), A88P (3/3), and A88T (4/4) did not contribute direct contacts (**Figure 4C**).

We then asked whether this apparent functional consistency with regard to interactions extended beyond SARS-CoV-2–specific antibodies. Using SAbDab, we analyzed mutated IGHV1-69 and IGHV3-30 antibodies targeting diverse foreign antigens, including viral, bacterial, protozoan, and allergen proteins. Remarkably, the same clonotype-shared SHMs were observed with broadly similar structural roles (**Figure 4B, C**). Among non–SARS-CoV-2 IGHV1-69 antibodies, S31N predominantly mediated antigen contacts (13/17), and Y95F consistently formed light chain contacts (21/21). A58T generally did not participate in contacts but formed antigen contacts in all human immunodeficiency virus 1 (HIV-1)-specific antibodies in which it was present, suggesting that its structural role may depend on antigenic context (**Figure 4B**). Among non–SARS-CoV-2 IGHV3-30 antibodies, V50L primarily mediated light chain contacts (7/8), whereas M34I (7/7), A88P (15/16), and A88T (6/6) largely lacked direct contact contributions. In contrast to SARS-CoV-2–specific antibodies, S31N less frequently contacted antigen (4/14), whereas K65R consistently mediated light-chain interactions (4/4) (**Figure 4C**).

Notably, all identified clonotype-shared SHMs in IGHV1-69 and IGHV3-30 arose within WRCY AID hotspot motifs, with the exception of F29L in IGHV1-69 and K65R in IGHV3-30 (**Figure 4B, C**), suggesting that the high frequency with which these mutations are acquired in an antigen-independent manner, may be driven, at least in part, by intrinsic sequence mutability.

Collectively, these findings indicate that clonotype-shared SHMs are not only recurrent but also frequently adopt conserved structural roles across antibodies targeting distinct epitopes and unrelated antigens. This recurrent behavior supports their classification as “pan-antigenic” SHMs, representing commonly accessible mutational solutions that recur across independent antibody responses.

### Clonotype-shared SHMs represent the most common substitutions at the respective antibody residue positions in healthy antibody repertoires

The presence of an AID hotspot and high intrinsic mutability does not necessarily determine which amino acid substitution will arise, because AID-mediated cytidine deamination can generate multiple nucleotide substitutions through downstream DNA repair pathways, and codon identity further influences the resulting amino acid change^29,30^. We therefore asked whether clonotype-shared SHMs occur at AID hotspots and represent the most frequently observed amino acid substitutions at their respective residue positions across the broader antibody repertoire.

To address this, we analyzed healthy, non-naïve BCR repertoires derived from peripheral blood (n = 3) and tonsillar tissue (n = 6) obtained from the Observed Antibody Space (OAS) database. We restricted the analysis to mutated (≥1 SHM) IGHV1-69 and IGHV3-30 sequences and calculated the frequency distribution of all observed amino acid substitutions at positions where clonotype-shared SHMs had been identified (**Figure S13A-O**). Although CDR positions generally exhibited substantial substitutional diversity, some residues showed highly constrained mutational profiles, including position 95 in IGHV1-69 (**Figure S13H**). Notably, at every position analyzed, the clonotype-shared SHM was the most frequently observed substitution across donors. Y95F was particularly striking, accounting for nearly all substitutions observed at heavy-chain position 95 in IGHV1-69 sequences (**Figure S13H**).

These findings indicate that clonotype-shared SHMs are not only recurrent in antigen-specific antibodies but also represent the dominant amino acid substitutions at their respective residue positions in polyclonal, antigen-agnostic repertoires from healthy individuals, consistent with strong intrinsic mutational accessibility and/or repertoire-level selection.

### Pan-antigenic SHMs rarely contribute to *de novo* contacts and are not sufficient for affinity maturation

Thus far, we defined clonotype-shared SHMs in the context of spike-specific GC BCR clonotypes using IGHV1-69 and IGHV3-30. However, analogous recurrent mutations may occur across other V genes and represent more broadly shared “pan-antigenic” SHMs. To generate a more general catalogue of such mutations, we analyzed antibodies in SAbDab encoded by the five most frequently represented heavy- and light-chain V genes (**Table S2**). For each V gene, we defined *pan-antigenic SHMs* as amino acid substitutions observed in at least 10% of antibodies using that gene (**Figure 5A**). This analysis identified numerous additional pan-antigenic SHMs beyond those previously characterized in spike-specific IGHV1-69 and IGHV3-30 antibodies (**Figure 5B, C**). Across all V genes examined, these mutations occurred in antibodies recognizing diverse antigens, further supporting their classification as pan-antigenic SHMs (**Figure 5B, C, S14A, B**).

**Figure 5:**
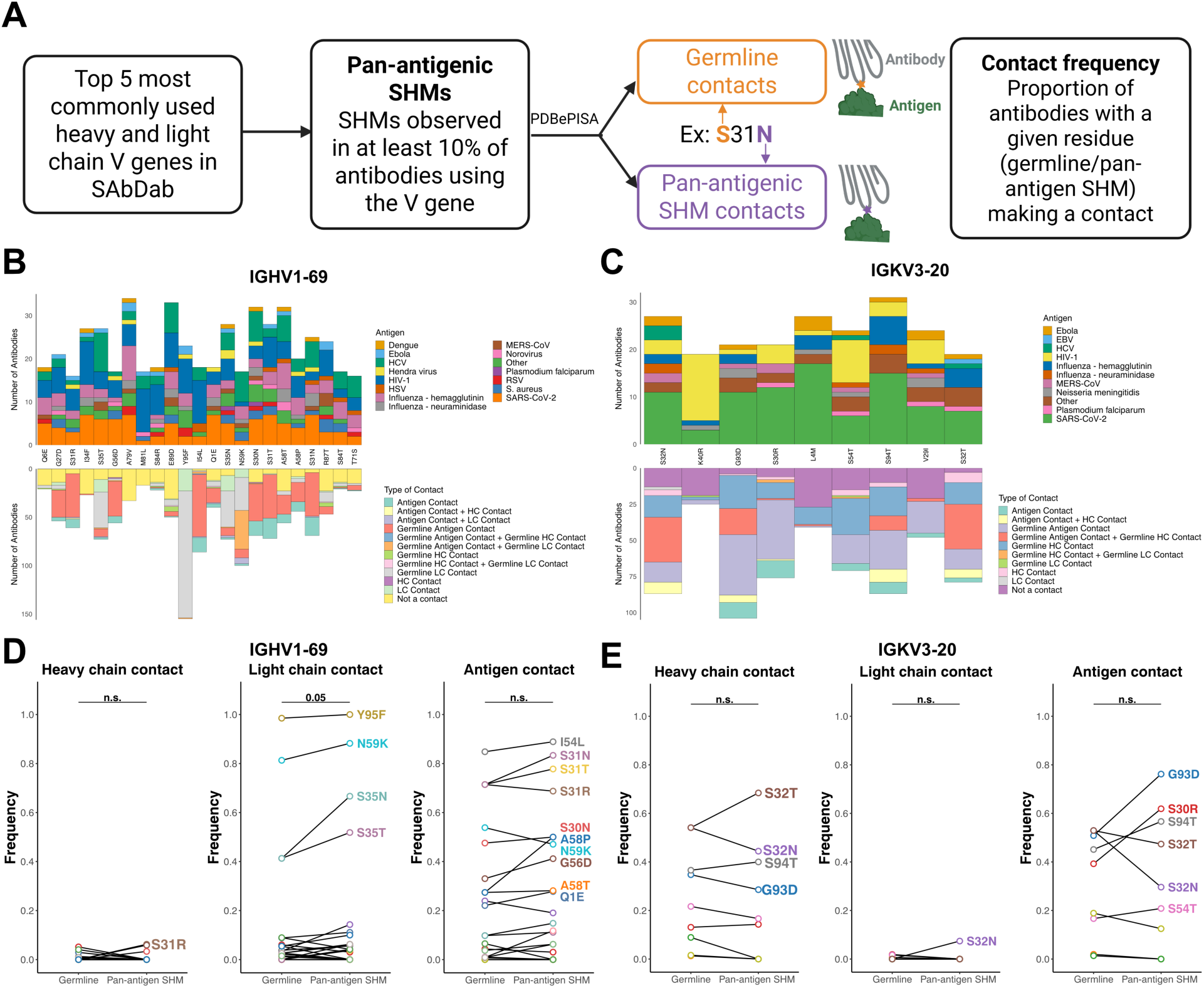
Contributions of pan-antigenic SHMs to antigen–antibody interactions. (**A**) Schematic representation of the approach used to define and compute germline and pan-antigenic SHM contact frequencies. (**B–C**) Distribution of antigen specificities recognized by antibodies harboring a given pan-antigenic SHM (top panels), and the types of contacts formed either by the mutated residue or by the corresponding germline residue when the mutation is absent (bottom panels), for pan-antigenic SHMs in (**B**) IGHV1-69 and (**C**) IGKV3-20. (**D–E**) Contact frequencies for pan-antigenic SHMs in (**D**) IGHV1-69 and (**E**) IGKV3-20. Contact frequency is defined as the proportion of antibodies in which a given residue, either mutated or germline, forms a contact with the heavy chain (left), light chain (center), or antigen (right). Linked dots represent paired comparisons of contact frequency for the same residue in its germline versus mutated (pan-antigenic SHM) state. Each color corresponds to a distinct pan-antigenic SHM. The non-parametric Wilcoxon signed rank test was used to compute p-values corresponding to pairwise comparisons between contact frequencies in (**D**) and (**E**).

Although some clonotype-shared SHMs contributed to structural contacts across antibodies of diverse antigen specificities, this observation alone does not establish that these mutations enhance affinity. To assess whether pan-antigenic SHMs preferentially create or preserve structural contacts, we compared the contact frequency of each mutation with that of the corresponding germline residue. We defined contact frequency as the proportion of antibodies in which the residue formed an antigen or interchain contact when present in its mutated or germline state. Although pan-antigenic SHMs frequently participated in antigen and/or cognate-chain contacts, the corresponding germline residues also commonly formed contacts (**Figure 5B, C, S14A, B, S15A, B**). Accordingly, most pan-antigenic SHMs showed no significant increase in contact frequency relative to germline (**Figure 5D, E, S15A, B**).

A small number of mutations represented notable exceptions, exhibiting substantial *de novo* contact formation, defined as >50% contact frequency for the mutated residue and with near-zero contact frequency for the corresponding germline residue. These included S35H and S35N in IGHV3-23 which formed light chain contacts; S59R, S57R, and G55S in IGHV1-46, which formed antigen and/or light chain contacts; Y32A and S53N in IGKV1-39 which formed antigen and/or heavy chain contacts, and S30R, S30G, and S93N in IGKV1-33 which formed antigen contacts (**Figure 5D, E, S15A, B**). Such cases, however, were uncommon.

If pan-antigenic SHMs represent broadly useful affinity-enhancing solutions, they should improve binding across diverse antibody backgrounds. To test this hypothesis, we focused on IGHV1- 69/IGKV3-20, the most commonly represented heavy-light chain pair in SAbDab, which occurred among antibodies recognizing 11 distinct antigens (**Figure 6B**). We first selected pan-antigenic SHMs that formed antigen or cognate-chain contacts in antibodies targeting at least five distinct antigens (“contact pan-antigenic SHMs”). Because non-contact mutations may also influence antibody structure or stability through epistatic interactions ^31,32^, we next performed linkage analysis between contact pan-antigenic SHMs and other SHMs in mutated IGHV1-69/IGKV3-20 sequences from healthy peripheral blood and tonsillar repertoires in Observed Antibody Space (**Figure 6A**). Several pan-antigenic SHMs showed significant linkage with additional heavy- and light-chain mutations, defined here as a linkage score ≥2 [log_2_(Linkage) ≥1] (**Figure S16A, B**). Restricting the analysis to linked mutations present in ≥10% of sequences identified two major non-contact pan-antigenic SHMs, A79V and E89D in IGHV1-69 (**Figure S16B**). Together, these analyses defined a minimal pan-antigenic SHM set comprising ten heavy-chain mutations (eight contact, two non-contact) and five light-chain mutations (all contact) for IGHV1-69/IGKV3-20 antibodies.

**Figure 6:**
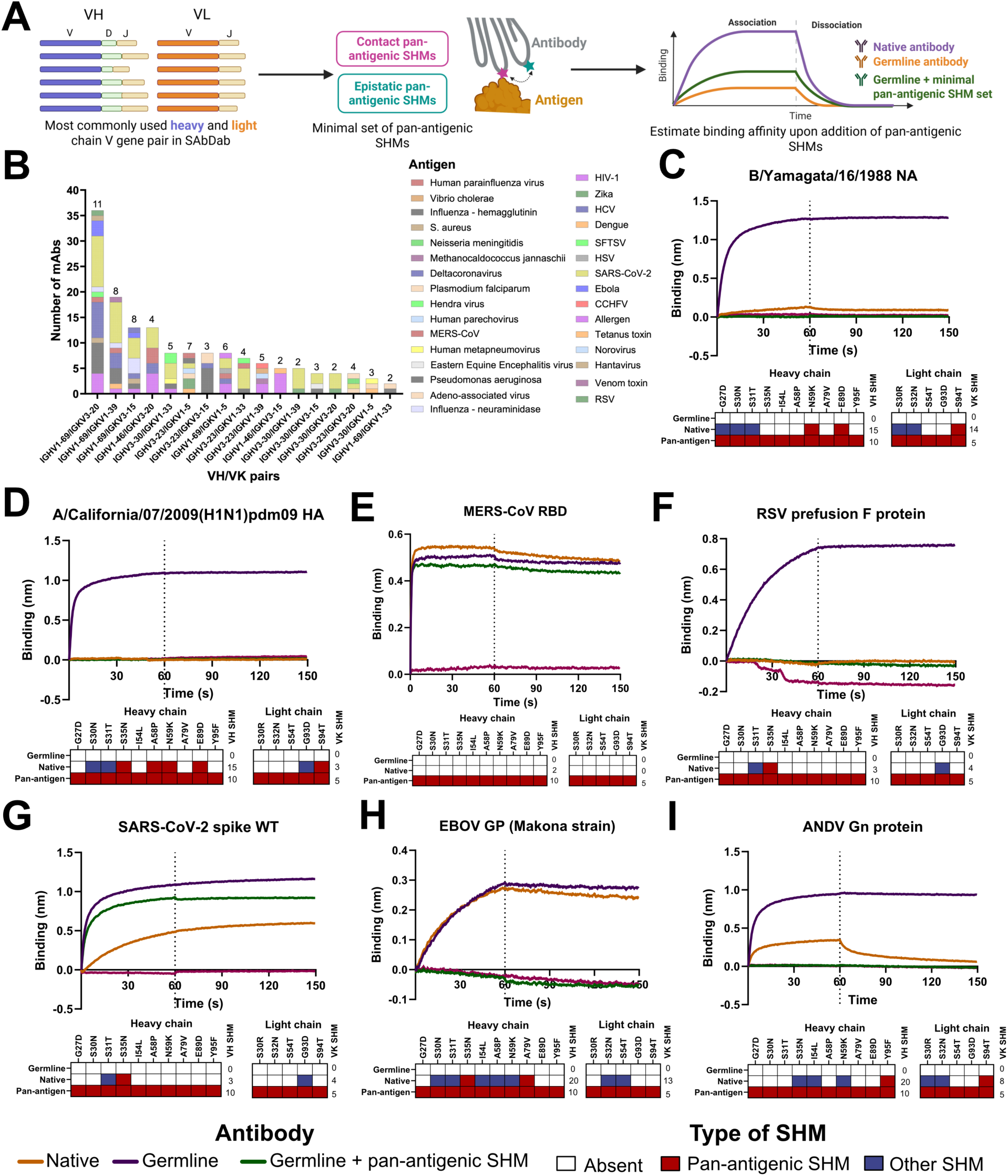
Contributions of pan-antigenic SHMs to affinity maturation across distinct germline backgrounds. (**A**) Schematic workflow illustrating the strategy used to identify contact and epistatically linked pan-antigenic SHMs for incorporation into different germline antibody backgrounds. (**B**) Frequency distribution of antigens recognized by antibodies using distinct heavy/light chain V gene pairs in SAbDab, ordered by decreasing abundance. The number above each bar indicates the total number of unique antigens recognized by antibodies using the corresponding V gene pair. (**C–G**) Biolayer interferometry (BLI) analysis of binding affinities for antibodies specific to (**C)** influenza B/Yamagata/16/1988 neuraminidase, (**D**) influenza A/California/07/2009 (H1N1)pdm09 hemagglutinin, (**E**) MERS-CoV RBD, (F) RSV prefusion F protein, and (**G**) SARS-CoV-2 wild-type spike protein. For each antigen, antibodies were expressed in three variants: native mature antibody from SAbDab (purple), germline-reverted antibody with all V gene mutations reverted (orange), and germline antibody incorporating the minimal pan-antigenic SHM set (green). Binding affinities of these variants were compared. The heatmaps below each panel indicate the mutational status at residues corresponding to the selected pan-antigenic SHMs: white denotes germline residue (no mutation), pink denotes the selected pan-antigenic SHM, and blue denotes a mutation distinct from the selected pan-antigenic SHM. The dotted vertical line separates association and dissociation curves in BLI.

To determine whether this minimal pan-antigenic SHM set could enhance affinity across diverse antibody backgrounds, we selected IGHV1-69/IGKV3-20 antibodies recognizing seven unrelated antigens: influenza B neuraminidase (B/Yamagata/16/1988 strain), influenza A H1 hemagglutinin (A/California/07/2009 strain), Middle East Respiratory Syndrome Coronavirus (MERS-CoV) RBD, SARS-CoV-2 spike, Respiratory Syncytial Virus (RSV) prefusion F protein, Ebola virus glycoprotein (EBOV GP) (Makona strain, mucin deleted) and Andes virus (ANDV) Gn glycoprotein. For each antibody, we generated three variants: (i) the mature antibody, (ii) a germline-reverted variant in which V gene SHMs were reverted, and (iii) the germline-reverted variant containing the minimal pan-antigenic SHM set. We measured binding to cognate antigens by biolayer interferometry. For most antibodies, introducing the pan-antigenic SHM set failed to enhance affinity relative to the germline-reverted variant (**Figure 6C-I**). The SARS-CoV-2– specific antibody DH1043, represented the sole exception, showing a modest increase in affinity after introduction of the pan-antigenic SHMs, although its affinity remained lower than that of the mature antibody (**Figure 6G**). Introduction of pan-antigenic SHMs however, resulted in loss of binding in antibodies against EBOV GP (**Figure 6H**) and ANDV Gn (**Figure 6I**).

Collectively, these findings indicate that pan-antigenic SHMs rarely generate *de novo* contacts and, as a group, are insufficient to drive affinity enhancement across diverse antibody backgrounds. Their recurrent acquisition may therefore reflect intrinsic mutational accessibility more than consistent positive selection to increase affinity. In contrast to clonotype-restricted SHMs, which appear to encode antigen- or epitope-specific solutions, pan-antigenic SHMs may represent broadly permissive substitutions that are repeatedly sampled during germinal center diversification and only occasionally contribute directly to affinity enhancement.

## DISCUSSION

Generally, antibody features at the genetic level fail to accurately encode information on the epitope targeted, thus serving as a major barrier towards modeling antigen antibody interactions. During affinity maturation, SHM generates sequence diversity upon which antigen-driven selection acts, allowing antibody lineages to explore mutational solutions compatible with recognition of their cognate epitopes. However, germinal center selection is inherently permissive, and a substantial fraction of acquired SHMs are affinity-neutral^17^. This background mutational variation limits the interpretability of individual substitutions when considered in isolation. Our results address this limitation by demonstrating that the clonotype specificity of SHMs can distinguish epitope-informative mutations from broadly recurrent mutations favored by intrinsic mutational accessibility. By comparing mutation frequencies across clonotypes using the same V gene, this framework helps disentangle the relative contributions of AID targeting biases and antigen-driven selection to observed SHM patterns. Affinity maturation can therefore provide not only a record of affinity optimization but also a readout of the structural and biochemical constraints imposed by the targeted epitope.

We identified and structurally characterized three antibodies, namely CR5, CR16 and CR18, derived from SARS-CoV-2 spike-specific GC B cells elicited by mRNA vaccination. All three antibodies harbored multiple clonotype-restricted SHMs and recognized cryptic epitopes on spike. CR5 and CR18 recognized a cryptic epitope on RBD that overlaps the site recognized by CR3022. an antibody previously characterized by Yuan and colleagues. They showed that the prefusion spike trimer sterically occludes this epitope and that adjacent RBD protomers must adopt the “up” conformation and undergo rotational rearrangements to permit antibody access^27^. These conformational changes relieve steric clashes between the CR3022 variable domain and neighboring RBD and S2 regions, as well as between the antibody constant region and the NTD. Subsequent structural studies demonstrated that CR3022 binding can destabilize the spike trimer and promote S1 subunit dissociation, further exposing the cryptic epitope^33^. The CR3022 light chain also contacts the NTD, underscoring the conformationally restricted nature of this binding mode. Although CR3022 was originally isolated from an individual infected with SARS-CoV, CR3022 fails to neutralize SARS-CoV-2, similar to CR5, because four amino acid differences within the cryptic epitope reduce its binding affinity^27,33,34^. More recently, Fu and colleagues showed that sequential immunization of mice harboring adoptively transferred CR3022-like B cells drove SHM-dependent expansion of neutralization breadth to include SARS-CoV-2 variants^24^. Together, these studies suggest that affinity maturation can refine antibodies targeting this cryptic epitope to expand their neutralization breadth.

Strikingly, all five clonotype-restricted SHMs in CR5 occurred in the IGLV3-25 light chain. This pattern parallels our previous observations for the public antibody M15, in which clonotype-restricted SHMs in the IGKV3-20 light chain contributed to recognition of a cryptic S2 epitope^12^. None of the clonotype-restricted SHMs in CR5 directly contacted antigen; instead, our structural and BLI analyses indicate that they reorganize the paratope and facilitate productive association with the cryptic epitope. These findings raise the possibility that light-chain diversification can play an important role in reshaping the paratope to accommodate sterically constrained or conformationally masked epitopes.

Several studies support a broader role for light-chain adaptation in antibody maturation. In VRC01-class antibodies targeting HIV-1 envelope protein, SHMs within the IGKV3-20 light chain stabilize the paratope and mitigate clashes with dynamic Env glycans^31^. The CAP256-VRC26 broadly neutralizing antibody lineage, which targets the V2 apex of HIV-1 Env, accumulates rare light chain mutations that reshape the paratope and enhance neutralization potency^35^. In the HyHEL-10 lysozyme-binding antibody family, a light chain mutation S30G alters binding thermodynamics and conformational flexibility without directly contacting antigen^36^. Similarly, in public IGHV3-53/66 class of SARS-CoV-2 RBD-directed antibodies, heavy chain SHMs refine interactions with conserved RBD residues, whereas alternative light chain pairings modulate shape complementarity and enable recognition of viral variants^37^. More broadly, the importance of non-contact epistatic interactions in reshaping the paratope is exemplified by a hapten-specific antibody in which SHM contributed to a ∼30,000-fold increase in affinity without introducing direct antigen contacts^32^.

Collectively, these observations highlight the capacity of light-chain sequence variation, through SHM or alternative pairing, to remodel paratope architecture without necessarily creating new antigen contacts. Such adaptations may be particularly important for antibodies recognizing cryptic, conformationally restricted, or evolutionarily variable epitopes. In this context, the concentration of clonotype-restricted SHMs within the light chain of CR5, together with our previous observations for M15^12^, suggests that light-chain maturation can provide epitope-specific structural solutions for accessing otherwise constrained antigenic surfaces.

Our structural analysis of CR16 identified a deeply buried NTD epitope that is almost entirely occluded in available prefusion spike conformations, raising the question of how B cells gain access to this otherwise inaccessible surface. To our knowledge, BLN8 represents the only other antibody reported to recognize this buried NTD epitope. Investigators isolated BLN8 from a coronavirus disease 2019 (COVID-19) convalescent donor^28,38^, indicating that immune responses can target this site following both infection and mRNA vaccination. One potential mechanism for exposing this epitope is dissociation or shedding of the S1 subunit, which occurs during spike-mediated membrane fusion and can also arise from spike expressed on host cells following mRNA vaccination. Consistent with this possibility, studies have detected circulating spike-derived antigens, including S1, following mRNA vaccination^39–42^. Thus, B cells may encounter conformationally altered or dissociated forms of S1 that expose NTD surfaces inaccessible within the intact prefusion spike trimer.

The enrichment of clonotype-restricted SHMs among spike-specific clonotypes that failed to bind spike by ELISA raises the possibility that some of these mutations arose under selection for alternative antigen specificities. However, these clonotypes may represent “dark” spike specificities directed toward cryptic or poorly accessible epitopes that standard *ex vivo* binding assays fail to detect because of limited sensitivity or antigen presentation^43^. Consistent with this possibility, six of the eighteen CR Fabs exhibited weak or undetectable binding, suggesting that at least some may recognize structurally occluded epitopes that are not captured under standard assay conditions. This observation is consistent with the broader association of clonotype-restricted SHMs and cryptic epitope recognition.

Analysis of CR3 clonotype further illustrated that increased SHM does not necessarily translate into improved antigen binding. A clonally related antibody that diverged from a common ancestor retained detectable spike binding by ELISA and biolayer interferometry, whereas the more highly mutated CR3 antibody did not. Thus, additional SHMs acquired within an evolving lineage can reduce, rather than enhance, measurable antigen binding. DeWitt and colleagues similarly showed that a substantial fraction of terminal “leaf” clones within antibody lineage trees display lower affinity relative to their inferred ancestors, potentially representing clones that have not yet been purged by affinity-based selection^44^. Thus, these observations emphasize the need to characterize clonotypes enriched for clonotype-restricted SHMs more deeply, particularly those with weak or undetectable antigen binding, and to determine whether such lineages are especially susceptible to SHMs that compromise affinity.

In contrast to clonotype-restricted SHMs, clonotype-shared SHMs recurred in antibodies targeting diverse antigens while using the same V gene. Our structural analyses further showed that these mutations often adopted conserved roles across independent antibodies, either contacting antigen or stabilizing heavy–light chain interactions. We therefore asked whether a minimal set of such pan-antigenic SHMs could enhance affinity across diverse antibody backgrounds and antigen specificities, which would suggest a degree of redundancy in the mutational solutions available during affinity maturation. Our experimental results did not support this model. Introducing a minimal pan-antigenic SHM set into diverse germline-reverted antibodies targeting seven unrelated antigens failed to improve affinity above germline levels, with a single SARS-CoV-2-specific antibody as the exception. Whether this isolated effect reflects structural features specific to its antigen-antibody interaction remains unclear. Overall, these findings argue against a model in which commonly sampled SHMs provide broadly transferable affinity gains. Instead, they support the view that affinity maturation remains predominantly epitope-tailored and depends on context-specific mutational solutions.

Given that many pan-antigenic SHMs arise at AID hotspot motifs, one might ask whether their recurrence simply reflects intrinsic mutational targeting rather than functional selection. These processes, however, operate at distinct levels. AID hotspot motifs influence where mutations arise, whereas downstream DNA repair, codon context, and selection determine which amino acid substitutions appear and persist within antibody repertoires. Thus, high mutational accessibility at a given position does not necessarily imply that the resulting amino acid substitution enhances antibody function. Consistent with this distinction, pan-antigenic SHMs rarely generated *de novo* contacts; that is, interactions absent from the corresponding germline residue. Instead, many produced chemically conservative substitutions that preserved existing structural interactions rather than creating new ones.

Importantly, many clonotype-restricted and clonotype-shared SHMs occur at the same residue positions yet produce different amino acid substitutions. Examples in IGHV1-69 include S35H versus S35N, F29M versus F29L, and A58I versus A58T. These patterns further demonstrate that the consequences of SHM depends not only on the position of mutation but also on the specific amino acid substitution, as differences in side-chain chemistry can differentially influence antigen engagement or inter-chain interactions. Previous studies support this principle. A large-scale analysis of SHMs in 196 antibodies revealed that substitution preferences are strongly dependent on both germline residue identity and sequence position^45^. Beem and colleagues incorporated this principle into ARMADiLLO, a computational framework that estimates the propensity for each possible amino acid substitution at individual positions within an antibody sequence^21^. Together, these observations indicate that recurrent mutational targeting can coexist with substantial diversity in amino acid outcomes, providing a substrate for epitope-specific adaptation during affinity maturation.

Overall, our findings support a model in which the extent to which a V gene SHM is shared across clonotypes provides information about its relationship to epitope recognition and affinity maturation. Clonotype-restricted SHMs frequently represent epitope-specific adaptations, particularly for structurally constrained or cryptic targets. In contrast, clonotype-shared SHMs recur broadly across antigen specificities and often exhibit pan-antigenic patterns, yet generally fail to enhance affinity across diverse germline backgrounds. Together, these results indicate that affinity maturation samples both highly specific, epitope-tailored mutational solutions and broadly accessible, largely permissive substitutions. Framing SHM along a spectrum of clonotype specificity provides a means to distinguish mutations that encode epitope-specific information from those shaped by intrinsic mutational accessibility, offering a framework for interpreting affinity maturation and informing antibody optimization.

## MATERIALS AND METHODS

### Clonotype-specificity analysis of SHMs among spike-specific germinal center B cells

#### Identifying SHMs in GC B cells

Paired heavy and light chain sequences from spike-specific GC B cells elicited upon SARS-CoV-2 mRNA vaccination were obtained from Kim et al., 2022 (https://zenodo.org/records/5895181).

Complete, productive and paired sequences derived from germinal center B cells were shortlisted. For the clonotype-specificity analysis, clone IDs assigned by Kim and colleagues were used as clonotype identifiers and only clonotypes with at least 20 clones were considered. Germline gene identities within each clonotype were re-verified by running the AssignGenes and ParseDb functions using the changeo package on Immcantation. Amino acid sequences of V genes of each clone were aligned to their corresponding germline V genes obtained from AssignGenes, using CLUSTALW, to identify somatic hypermutations.

#### Computing clonotype-specificity of SHMs

Frequency of each SHM within a clonotype was computed as the number of clones possessing the SHM in the clonotype. Mutations occurring in at least 30 percent of clones within a clonotype were considered as ‘frequent mutations’. Clonotypes were grouped based on their V gene usage (heavy and light, separately). Heavy and light chain V genes used by at least 10 clonotypes and presenting at least 10 frequent mutations were then shortlisted for further analysis.

For each frequent SHM, the number of clonotypes presenting the SHM frequently (at least 30 percent of clones) using the given V gene, was used as an estimate of clonotype-specificity, depicted in Figure 1 – higher the sharing, lower the clonotype-specificity. Clonotype-shared SHMs were defined as SHMs shared frequently across at least five clonotypes using the given V gene. Clonotype-restricted SHMs were defined as SHMs exclusive to a clonotypes using the given V gene.

### Selecting clones with clonotype-restricted SHMs for assessing epitope-specificity

Clonotypes with at least three clonotype-restricted SHMs (heavy or light chain V genes) were shortlisted and further filtered based on whether the clonotype was assigned as Spike-positive on ELISA by Kim et al. The most mutated clone with all clonotype-restricted SHMs reported for the clonotype was selected for expression as a monoclonal Fab for each clonotype (labelled clonotype-restricted (CR) Fabs). For cases where the clonotype-restricted SHMs were split across sub-lineages, the most mutated clone within each sub-lineage with the clonotype-restricted SHMs was selected (CR15-16 and CR17-18).

### Monoclonal Fab expression and purification

The VH and VK genes of each Fab were cloned into pcDNA3.4 mammalian expression vectors (GenScript). Expi293 cells (25- to 200-mL cultures) were transfected with the heavy- and light-chain-expressing vectors at a 1:1 ratio by concentration, along with Expifectamine (transfection reagent; Thermo Fisher Scientific), following the protocol described in the lipid-based Thermo Fisher Expi293™ Expression System Kit. The cells were incubated in a shaker flask at 37°C for 18-22 hours at 125 rpm. Enhancers 1 and 2 were added to the culture the next day, followed by further incubation for 5-6 days. Cells were then pelleted down by centrifugation and filtered (0.45 µm). The supernatant was mixed with Nickel-NTA agarose resin (Gold Biotechnology) (3-mL per 50-mL of supernatant) in a rotary shaker overnight at 4°C and then loaded onto Poly-Prep chromatography columns (Bio-Rad Laboratories, Inc.). Columns were washed with 10mM HEPES (pH 7.5), 150mM NaCl, 15mM Imidazole and eluted with 10mM HEPES (pH 7.5), 150mM NaCl, 400mM Imidazole. Purified Fabs were concentrated and stored at 4°C.

### ELISA and epitope binning

96-well Immulon 4 HBX plates (Thermo Scientific) were coated with 50 µL of the recombinant protein at 2 µg/mL and were stored at 4°C overnight. Wells were then washed three times with phosphate-buffered saline (3) containing 0.1% Tween-20 (PBST) and blocked with 100 µL PBST supplemented with 3% milk for 1 hour at room temperature. For ELISA with Fab, 100µL of antibody at 20µg/mL was added to each well in triplicates and incubated for 2 hours at room temperature. For ELISA with full-length IgG, antibodies were diluted 3-fold starting at 30µg/mL and 100µL of serially diluted antibodies were added to each well in duplicates and incubated for 1 hour at room temperature. For epitope binning, 50µL of the reference antibody at 10µg/mL and 50µL of the CR Fab at 200µg/mL were mixed and added to each well, followed by incubation for 2 hours at room temperature. As a positive control for competition, the biotinylated reference Fab at 10µg/mL was competed against its non-biotinylated version at 200µg/mL. As a negative control for competition, the biotinylated reference Fab at 10µg/mL was mixed with 1% milk (blank). The wells were then washed with PBST two times. Next, for ELISA with Fab, 100 µL of mouse anti-human Fab-specific antibody conjugated to horseradish peroxidase (Sigma, A3682) at a 1:3000 dilution was added to each well, followed by incubation for 1 hour at room temperature. For ELISA with full-length IgG, 100 µL of mouse anti-human IgG-specific antibody conjugated to horseradish peroxidase (Sigma, A56856) at a 1:3000 dilution was added to each well, followed by incubation for 1 hour at room temperature. For epitope binning, 100uL of streptavidin conjugated to horseradish peroxidase (Sigma, SA10001) at a 1:1000 dilution was added to each well, followed by incubation for 1 hour at room temperature. Plates were washed three times with PBST and developed by adding 100 µL of o-phenylenediamine dihydrochloride (OPD; Sigma-Aldrich) in phosphate citrate buffer (PCB; Sigma-Aldrich) with 0.04% hydrogen peroxide for 10 minutes. The reaction was stopped by adding 50 µL of 3 M hydrochloric acid (HCl; Fisher). The optical density (OD) was read for each well using a Synergy 4 (BioTek) plate reader at a wavelength of 490 nm. For ELISA, the average OD across triplicates or duplicates subtracted by the average of blanks was reported as the binding signal. For epitope binning, competition between a CR Fab and a reference Fab was quantified as the percentage decrease in mean OD across replicates, relative to the reference Fab alone (negative control).

### Live-virus neutralization assay

Microneutralization assays were performed as previously detailed^46^. In brief, Vero E6 TMPRSS2 cells were seeded in 96-well tissue culture plates (Corning, 3340) at a density of 1 × 10^4^ cells per well. After 24 hours, mAbs were diluted to an initial concentration of 30 µg/mL in 1x MEM with 2% FBS and then serially diluted 1:3 to achieve a final concentration of 0.041 µg/mL. An 80-µL portion of the mAb dilution was mixed with 80 µL of SARS-CoV-2 diluted to 10^4^ TCID_50_/mL, and this mixture was incubated at room temperature for 1 hour. Following incubation, 120 μL of the virus/mAb mixture was used to infect the cells for 1 hour. The inoculum was subsequently removed and replaced with 100 µL of 1x MEM containing 2% FBS and 100 µL of antibody dilution. The cells were then incubated at 37°C in a 5% CO_2_ environment for 48 hours before fixation with 10% paraformaldehyde (Polysciences) for 24 hours. After fixation, the paraformaldehyde was discarded, and cells were permeabilized by adding 100 µL of PBS containing 0.1% Triton X-100 (Fisher) for 15 minutes at room temperature. Once permeabilization was complete, the Triton X-100 solution was removed, and cells were blocked with 100 µL of 3% non-fat milk (Life Technologies) diluted in PBS. Finally, cells were stained for SARS-CoV-2 nucleoprotein using mAb 17C7, as previously described^46^.

### Cryo-EM sample and grid preparation

SARS-CoV-2 spike RBD was incubated with CR5 or CR18, RBD Class-1 and Class-5 Fabs at 1 mg/mL at a molar ratio of 1:1.2:1.2:1.2 RBD:CR5:Class-1:Class-5 for 30 minutes at 4°C. SARS-CoV-2 spike NTD was incubated with CR16 and PH017 at 1 mg/mL at a molar ratio of 1:1.4:1.4 NTD:CR16:PH017 for 30 minutes at 4°C. Immediately before grid preparation, fluorinated octyl maltoside was added to the pre-formed complex at 0.02% w/v final concentration. 3µL of the pre-mixed sample were applied on glow discharged (15 mA, 25 s) Quantifoil Cu R1.2/1.3 or UltrAuFoil 1.2/1.3 300 Mesh grids (Ted Pella Inc) and subsequently blotted for 4 to 5.5 seconds at blot force 1 under chamber conditions maintained at 4°C and 100% humidity and then plunge-frozen in liquid ethane using an FEI Vitrobot Mark IV (Thermo Fisher Scientific).

All datasets reported were collected at MSSM cryo-EM Core on a Glacios™ G2 (Thermo Fisher Scientific) operated at 200 kV and equipped with a X-FEG electron gun and Falcon IVi detector. Collection was done in counting mode with a cumulative dose of 50e^-^/Å^2^ at a magnification of 190,000, corresponding to a calibrated pixel size of 0.722 Å and a defocus range from -1.2 to - 2.4 µm.

### Cryo-EM data collection and processing

#### CR5-RBD Complex

368 and 10,386 movies were collected from a single grid. Movies were aligned and dose weighted using patch motion correction implemented in cryoSPARC V5.0.3^47^. Contrast transfer function estimation was done in cryoSPARC V.5.0.3 using Patch CTF. After micrograph denoising and blob picking, particles were extracted at a box size of 384 pixels with a 2x bin and subjected to a first round of 2D classification. 2D classes corresponding to multiple projections of the particles were selected and used for template picking (2,421,230 particles). The picked particles were extracted with a 3x binning and subjected to a 2D curation yielding 1,545,382 used to generate 3 initial models. The particles originating the best volume (∼34%) underwent 3D non-uniform refinement followed by one round of 3D heterogeneous refinement. Particle stack associated with the best occupancies and overall map quality were extracted at full box size and subjected to 3D non-uniform refinement (NUR), local and global CTF refinement and local refinement with a mask excluding the volume corresponding to the constant region of the Fab. 446,194 particles yielded a map at a nominal resolution of 3.66Å based on the gold-standard Fourier shell correlation od 0.143 criterion.

#### CR16-NTD Complex

506 and 10,115 movies were collected from a single grid. Movies were aligned and dose weighted using patch motion correction implemented in cryoSPARC v5.0.3^47^. Contrast transfer function estimation was done in cryoSPARC v.5.0.3 using Patch CTF. After micrograph denoising and blob picking, particles were extracted at a box size of 384 pixels with a 2x bin and subjected to a first round of 2D classification. 2D classes corresponding to multiple projection of the particles were selected and used for template picking (635,930 particles). The picked particles were used to generate 3 initial model followed by three rounds of 3D NUR and 3D heterogeneous refinement where only the best volume was kept for the subsequent cycle. This particle curation yielded a stack of 270,504 particles (∼43%) that underwent 3D NUR, local and global CTF refinement and local refinement with a mask excluding. Because of map anisotropy, the dataset was curated once more following the advanced automated workflow from Cryosparc^48^. The initial and curated volumes were used as template for 3D heterogeneous refinement from the whole initial particle set. The particles corresponding to the best volume were rebalanced to filter out particles in over-represented viewing directions, extracted to 1.5x binning and 3 initial models were built followed by 3D heterogeneous reconstruction. After 3D NUR reconstruction of a volume corresponding to the remaining 171,175, particles were rebalanced once more based on their orientation. The remaining 137,709 underwent 3D NUR, local and global CTF refinement and local refinement with a mask excluding the constant region of the Fabs, yielded a map at a nominal resolution of 4.07Å based on the gold-standard Fourier shell correlation od 0.143 criterion. Finally the map was subjected to anisotropy correction^49^.

#### CR18-RBD Complex

307 movies were collected. Movies were aligned and dose-weighted using cryoSPARC v5.0.3^47^. Contrast transfer function estimation was done in cryoSPARC V5.0.3^47^ using Patch CTF, and particles were picked with cryoSPARC’s blob picker. The picked particles were extracted with a box size of 384 pixels, with 3x binning, subjected to a 2D classification. 16,436 particles were used for reconstruction of a map at 9Å nominal resolution.

### Model Building and Refinement

The DeepEMhancer sharpened and anisotropy corrected focused maps for the CR5-RBD and CR16-NTD complexes respectively, were used for model building. PDB 6XDG was used as a starting model for RBD domain and REGN10933, 8S6M was used as starting model for S2H97 and an unpublished model was used as starting model for NTD and PH017. Individual CR Fab models were generated using AlphaFold3 server^50^ and rigid body fitted in the corresponding densities. The models were then subjected to multiple rounds of real-space refinement in Phenix^51^ and manual adjustment using Coot^52^. Finally, models were validated with MolProbity^53^. Figures and structural analysis were performed using ChimeraX^54^. The structural biology software was compiled and distributed by SBGrid.

### Analysis of antigen-antibody structures from SAbDab

Metadata for antigen-antibody structures from SAbDab were downloaded on 14^th^ July 2025^23^. FASTA sequences of the heavy and light chains for each structure were extracted from their PDB IDs using pdb-tools^55^. V genes for each FASTA sequence were then annotated using igblastp^56^. Structures with antibodies using the top 5 most common V genes in SAbDab (IGHV1-69, IGHV3-30, IGHV1-46, IGHV3-23, IGKV3-20, IGKV3-20, IGKV1-33, IGKV1-5, IGKV3-15, IGKV1-39) were then shortlisted. IGHV1-2, which was also among the top five most commonly used V genes, was excluded from the analysis due to a high bias towards HIV-1-specific antibodies. Next, antibodies against pathogens including viruses, bacteria, protozoa were shortlisted for further analysis.

Mutations in the V gene for each antibody were identified by aligning the V gene amino acid sequences of the heavy/light chains to the corresponding germline V gene sequences. Pan-antigenic mutations were then identified as those appearing in at least 10 percent of antibodies using the V gene in SAbDab.

### Residue contact analysis

Contacts between heavy chain, light chain, and antigen for each PDB structure were identified using PDBePISA^57^. Interfaces corresponding to the heavy and light chain identifiers annotated in SAbDab were extracted from the PDBePISA output. For each structure, intrachain (within heavy or light chain) and interchain (heavy–light or antibody–antigen) contacts were identified. Residues harboring pan-antigenic SHMs that formed contacts were designated as pan-antigenic contacts. Conversely, when the corresponding germline residue was present at the same position (i.e., the SHM was absent) and formed contacts, these were designated as germline contacts. Contact frequency was defined as the proportion of antibodies using a given V gene in which the residue, either mutated (pan-antigenic SHM) or germline, formed a contact, calculated relative to the total number of antibodies analyzed for that V gene.

### Repertoire analysis from healthy participants

#### Clonotype-shared SHM analysis

Residue positions corresponding to clonotype-shared SHMs identified in IGHV1-69 and IGHV3-30 from the spike-specific germinal center repertoire were first defined. Non-naïve BCR sequences harboring at least one SHM were retrieved from peripheral blood (n = 3) and tonsillar tissue (n = 6) of healthy donors from the Observed Antibody Space (OAS) database. Sequences annotated as using IGHV1-69 or IGHV3-30 were shortlisted for analysis. For each sequence, the V gene amino acid sequence was aligned to its assigned germline V allele as provided by OAS to identify substitutions at positions corresponding to the clonotype-shared SHMs. Sequences in which the germline allele encoded a different amino acid at the position of interest (i.e., not matching the reference residue associated with the clonotype-shared SHM) were excluded from analysis. All observed substitutions from the germline residue at these positions were catalogued, and their frequencies were computed within each donor’s repertoire for comparative analysis.

#### Linkage analysis

Non-naïve BCR sequences using IGHV1-69 and IGKV3-20 were shortlisted and pooled across the nine donors for the pan-antigenic SHM linkage analysis. Linkage between each pan-antigenic SHM identified in IGJV1-69 and IGKV3-20 and every other SHM in the repertoire was computed as the ratio of the frequency of sequences presenting both SHMs to the product of frequencies of the individual mutations.

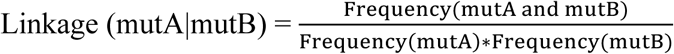

### Selection of pan-antigenic SHMs for affinity analysis

Two types of pan-antigenic SHMs in IGHV1-69 and IGKV3-20 genes were selected for introduction into different germline backgrounds:

#### Contact pan-antigenic SHMs

Pan-antigenic SHMs were defined as mutations that formed at least one type of contact (intra- or inter-chain) across antibodies targeting at least five distinct antigens. In cases of ties, SHMs were prioritized based first on multifunctionality (i.e., the ability to form contacts with multiple chains), and second on their overall frequency of occurrence across antigens in SAbDab.

#### Epistatic pan-antigenic SHMs

For identification of epistatic SHMs, linkage was evaluated only between contact pan-antigenic SHMs and mutations observed in ≥10% of sequences. Mutations with linkage >1 to at least one contact pan-antigenic SHM were designated as epistatic pan-antigenic SHMs.

### Biolayer interferometry (BLI)

Anti-human IgG Fc biosensors (Sartorius; 18-5001) were hydrated in 1X PBS, 0.1% Tween and 0.2% Bovine Serum Albumin buffer for 10 minutes. Then, 10 µg/mL of full-length IgG antibodies diluted in the same buffer was immobilized on a biosensor, and purified recombinant protein was flowed at 10µM, using the ForteBio Sartorius BLItz System. The steps involved are briefly described as follows: i) Baseline: the biosensor was initialized at baseline by dipping it in buffer for 30 seconds, ii) Loading: 4 µL of the antibody (M15) at 10 µg/mL was loaded onto the biosensor for 90 seconds, iii) Baseline: the biosensor was dipped back in buffer for 30 seconds, iv) Association: 4 µL of the antigen at 10µM was loaded onto the biosensor for 60 seconds, and v) Dissociation: the biosensor was dipped back into the buffer to let the antigen dissociate for 90 seconds. A control run with buffer loaded instead of the antibody and a negative control run with an irrelevant Mpox A35-specific antibody (EV35-2) were performed to confirm specific binding^58^. Binding signal with the reference control run subtracted was plotted over association and dissociation and compared across the native, germline and germline with pan-antigenic SHM antibodies.

## Acknowledgements

We thank Dr. Julian Zhou and Dr. Ali H. Ellebedy at Washington University in St. Louis for generously sharing information on the GC BCR sequences that were used in this study. We acknowledge Dr. Gagandeep Singh, Dr. Jordan J. Clark and Disha Bhavsar from the Krammer laboratory at the Icahn School of Medicine at Mount Sinai, Dr. Jason McLellan from University of Texas, Austin, Dr. Charles Daniel Murin and Dr. Andrew Ward from Scripps Research Institute for providing viral glycoprotein constructs that were critical for the study. Schematics for figures were created with BioRender.com and Affinity Designer.

## Funding

National Institutes of Health grant R01 AI168178 (GB)

National Institutes of Health grant FIRST U54CA267776 (CHC)

National Institutes of Health grant 1S10OD036213-01 (Cryo-EM CoRE, Icahn School of Medicine at Mount Sinai)

Irma T. Hirschl/Monique Weill-Caulier Trust (GB)

National Center for Advancing Translational Sciences Clinical and Translational Science Awards (CTSA) grant UL1TR004419 (Scientific Computing and Data at the Icahn School of Medicine at Mount Sinai)

National Institutes of Health grant S10OD026880 (GB)

National Institutes of Health grant S10OD030463 (GB)

National Institutes of Health grant G20AI174733 (BSL-3 Biocontainment CoRE)

## Author contributions

Conceptualization: VNR, CHC, GB

Methodology: VNR, CHC, GB

Investigation: VNR, BB, AC, AA

Visualization: VNR, BB, GB

Funding acquisition: CHC, GB

Supervision: CHC, GB

Writing – original draft: VR, BB, GB

Writing – review & editing: VR, BB, AA, FK, CHC, GB

## Data and Materials Availability

Paired heavy and light chain sequences from COVID-19 mRNA vaccinee spike-specific GC B cells were obtained from Kim et al., 2022 (https://zenodo.org/records/5895181). The EM maps have been deposited in the Electron Microscopy Data Bank (EMDB) under accession code EMD-78683 (CR5 complex), EMD-78684 (CR16 complex) and EMD-78896 (CR18 complex), and the accompanying atomic coordinates in the Protein Data Bank (PDB) under accession code 38AN (CR5) and 38AO (CR16). Additional data supporting the findings can be found in the supplemental information. All reagents will be made available upon request after completion of a ‘Materials Transfer Agreement’.

