## Supplementary Material for "Clonotype specificity distinguishes epitope-tailored and permissive somatic hypermutation"

### Supplementary File

#### Tables

| | **CR5-RBD-S2H97-REGN10933** (EMD-78683) (PDB 38AN) | **CR16-NTD-PH017** (EMD-78684) (PDB 38AO) |
|---|---|---|
| **Data collection and processing** | | |
| Magnification | 190,000 | 190,000 |
| Voltage (kV) | 200 | 200 |
| Electron exposure (e–/Å$^2$) | ~50.0 | ~50.0 |
| Defocus range (μm) | -1.2, -2.4 | -1.2, -2.4 |
| Pixel size (Å) | 0.722 | 0.92 |
| Symmetry imposed | C1 | C1 |
| Initial particle images (no.) | 1,545,382 | 1,541,275 |
| Final particle images (no.) | 44619 | 137,709 |
| Map resolution (Å) | 3.66 | 4.07 |
| FSC threshold | 0.143 | 0.143 |
| Map resolution range (Å) | 3.0-4.0 | 3.4-5.0 |
| **Refinement** | | |
| Initial model used (PDB code) | | |
| Model resolution (Å) | 3.7 | 4.1 |
| FSC threshold | 0.143 | 0.143 |
| Map sharpening *B* factor (Å$^2$) | 182.2 | 131.6 |
| Model composition | | |
| Non-hydrogen atoms | 6887 | 5440 |
| Protein residues | 891 | 690 |
| Nucleotide | --- | --- |
| Ligands | --- | --- |
| *B* factors (Å$^2$) | | |
| Protein | 39.58 | 29.92 |
| Nucleotide | --- | --- |
| Ligand | --- | --- |
| R.m.s. deviations | | |
| Bond lengths (Å) | 0.004 | 0.003 |
| Bond angles (°) | 0.626 | 0.680 |
| Validation | | |
| MolProbity score | 0.97 | 1.11 |
| Clashscore | 0.81 | 1.58 |
| Poor rotamers (%) | 0.00 | 0.00 |
| Ramachandran plot | | |
| Favored (%) | 96.58 | 96.70 |
| Allowed (%) | 3.42 | 3.30 |
| Disallowed (%) | 0.00 | 0.00 |

**Table S1: Cryo-EM data collection, refinement and validation statistics**

| Heavy chain V gene | Number of antibodies | Light chain V gene | Number of antibodies |
|---|---|---|---|
| IGHV1-69 | 158 | IGKV3-20 | 177 |
| IGHV3-30 | 105 | IGKV1-33 | 72 |
| IGHV3-23 | 83 | IGKV1-39 | 124 |
| IGHV1-46 | 46 | IGKV1-5 | 69 |
| IGHV1-2[#] | 87 | IGKV3-15 | 73 |

**Table S2: Top five most commonly used heavy and light V genes in SAbDab, used for pan-antigenic SHM analysis.** [#] IGHV1-2 was removed from further analysis due to extensive representation of HIV-1 specific antibodies in the database, which could potentially create biases in the pan-antigenic SHM analysis.

### Figures

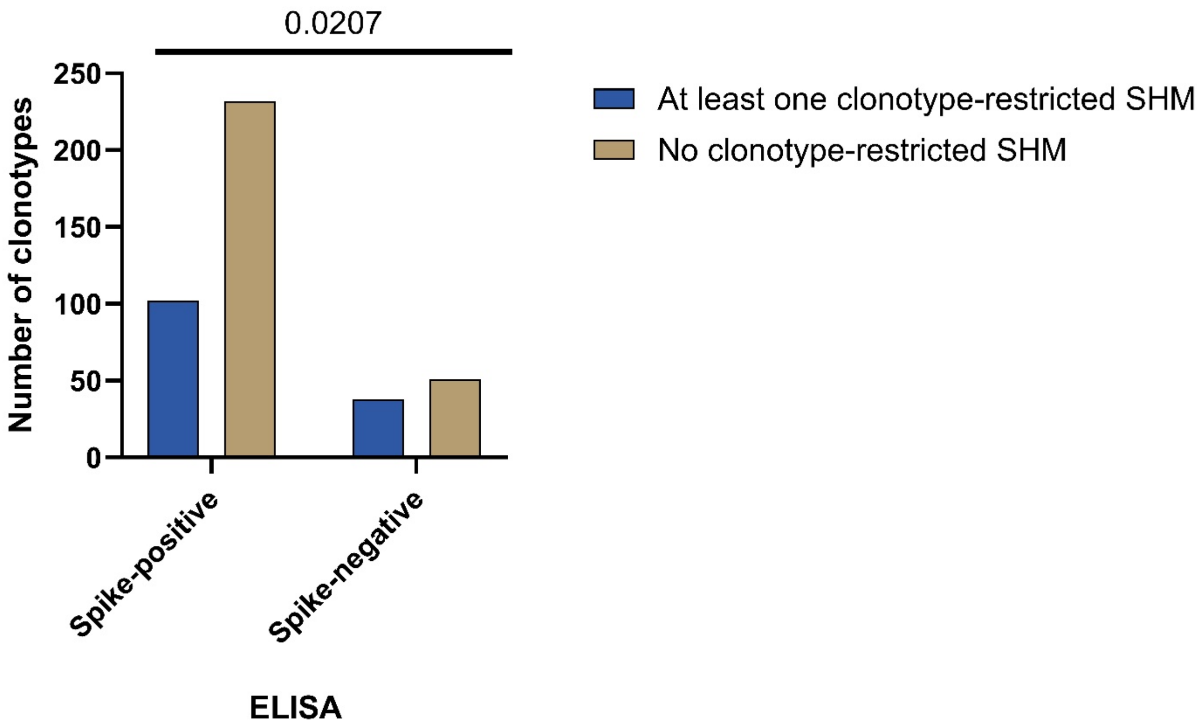

**Figure S1: Occurrence of clonotype-restricted SHMs in spike-positive and spike-negative clonotypes on ELISA.** Bar plot representing the number of spike-positive and spike-negative clonotypes that possess no clonotype-restricted SHMs (beige) versus at least one clonotype-restricted SHM (blue). Chi-squared ($\chi^2$) contingency test was used to test statistical significance for association.

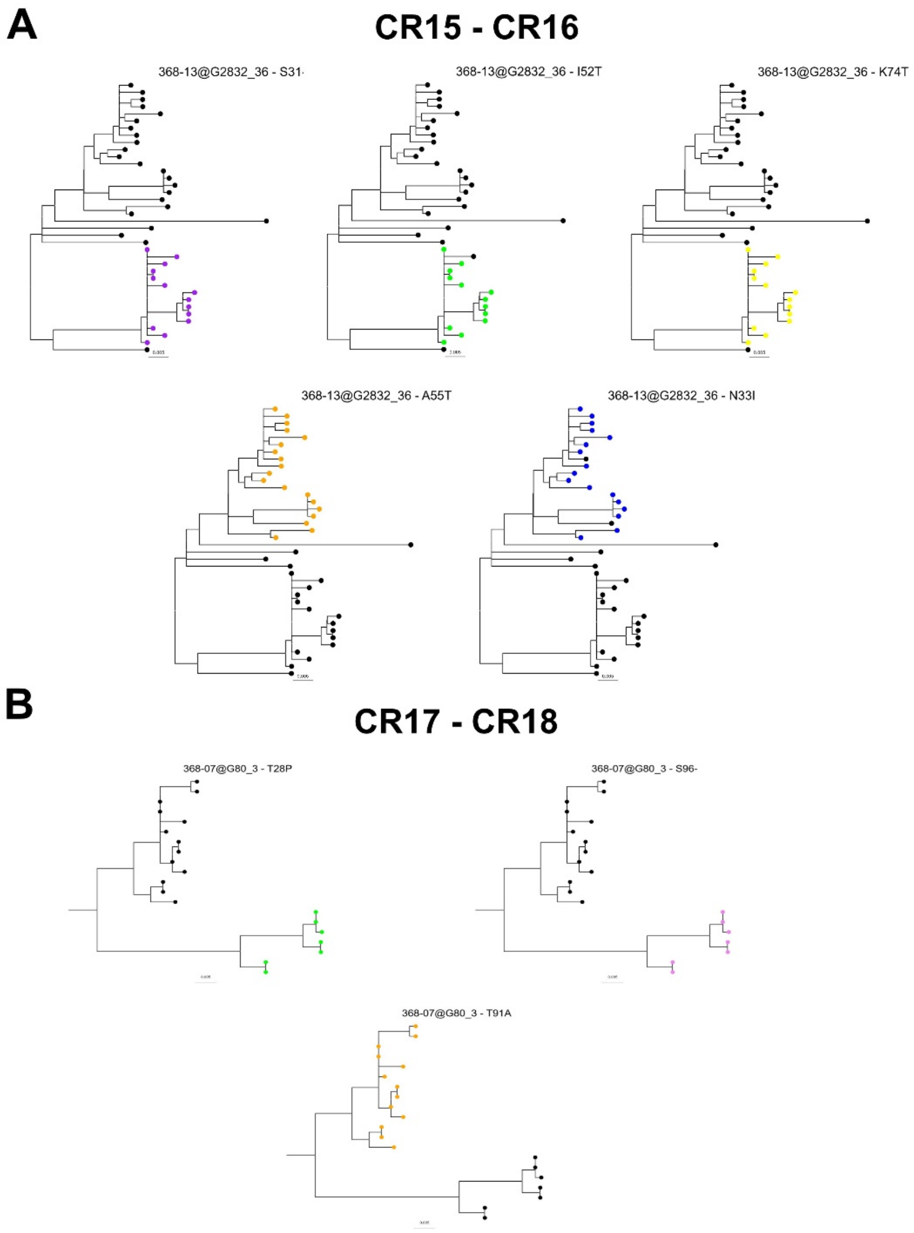

**Figure S2: Distribution of clonotype-restricted SHMs in clonal lineages corresponding to CR15-18. (A and B)** Phylogenetic reconstruction of clonal lineage trees of clonotypes harboring **(A)** CR15-16 and **(B)** CR17-18 clones, representing the distribution of each clonotype-restricted SHM observed in these clonotypes. Each tip represents an observed clone, and colored tips represent clones that possess the clonotype-restricted SHM. Scale for branch length is provided below each tree.

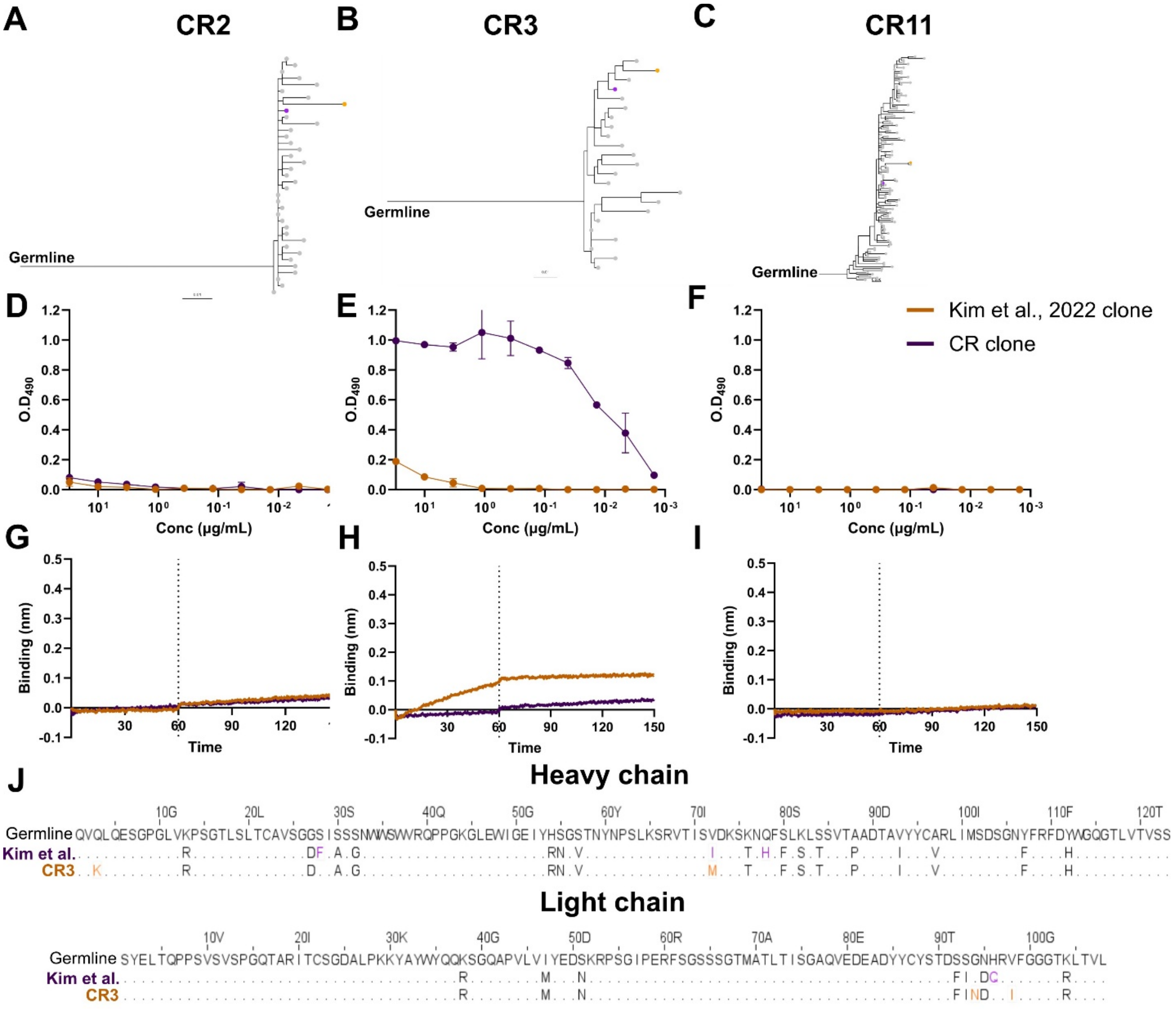

**Figure S3: Full-length IgG binding analysis of CR Fabs with undetectable binding on Fab ELISA.**
**(A-C)** Clonal lineage inference of spike-specific GC clonotypes bearing **(A)** CR2, **(B)** CR3 and **(C)** CR11 reconstructed using Cloanalyst. The clone chosen by Kim and colleagues is colored orange while the CR clone is colored purple.
**(D-F)** ELISA curves of full-length IgG **(D)** CR2, **(E)** CR3 and **(F)** CR11 antibodies against spike protein. The x-axis is represented in inverted log-scale.
**(G-I)** BLI analysis of binding affinities of **(G)** CR2, **(H)** CR3 and **(I)** CR11 to spike protein. The dotted vertical line separates association and dissociation curves in BLI.
**(J)** Sequence alignment of the heavy (above) and light chain (below) variable regions of CR3 and its counterpart from Kim and colleagues. Mutations exclusive to the clone from Kim et al are colored in purple while those exclusive to CR3 are colored in orange.

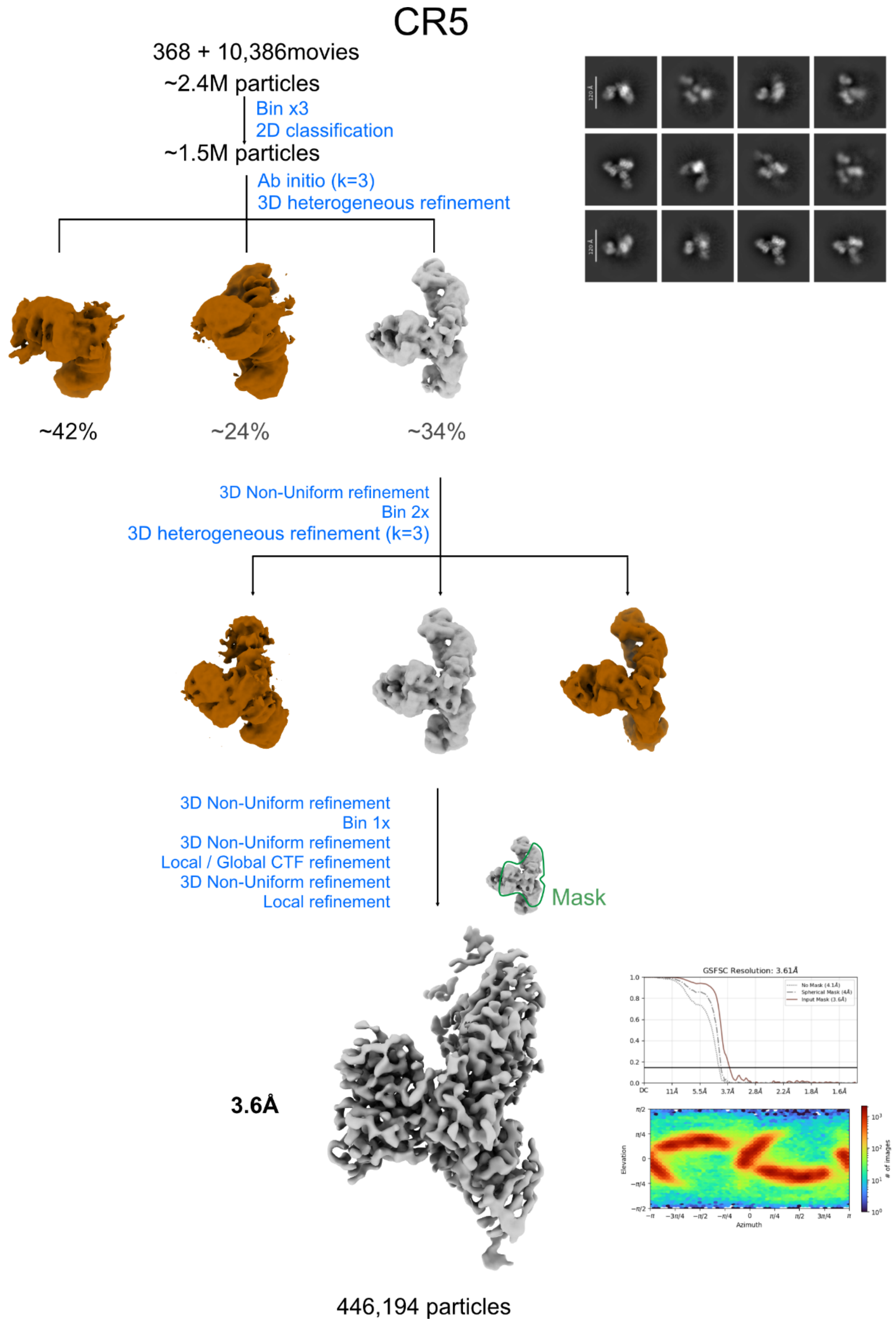

**Figure S4. Cryo-EM sorting and image processing workflow for SARS-CoV-S RBD in complex with CR5, S2H97 and REGN10933.**

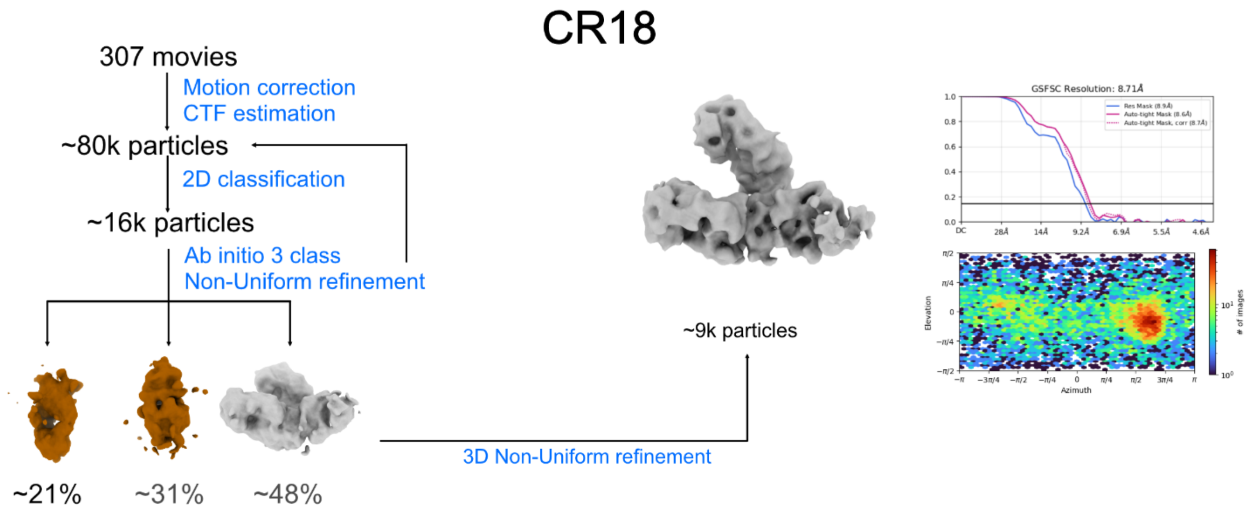

**Figure S5. Cryo-EM sorting and image processing workflow for SARS-CoV-S RBD in complex with CR18, S2H97 and REGN10933.**

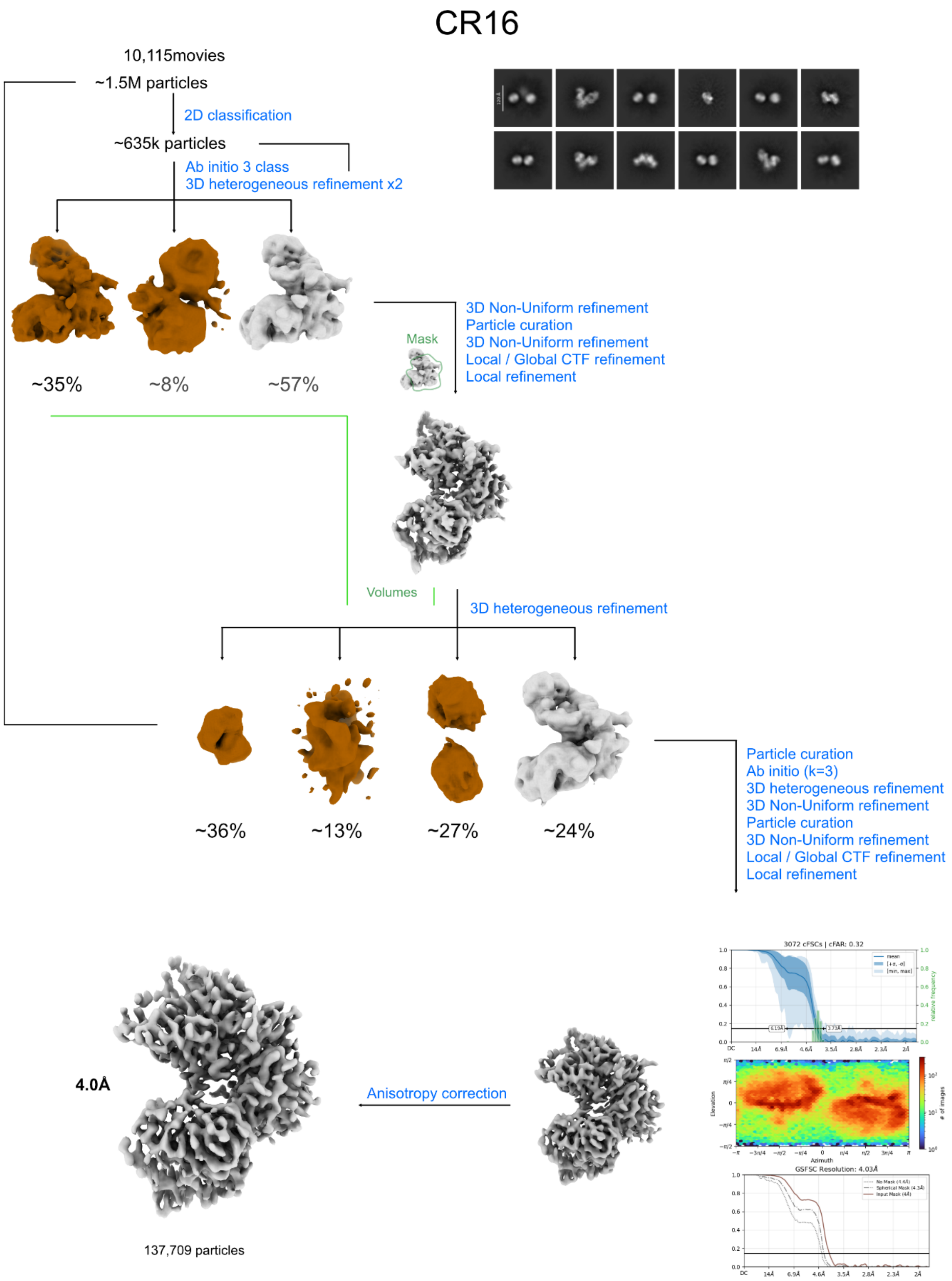

**Figure S6. Cryogenic electron microscopy data processing workflow for CR16-NTD-PH017 complex.**

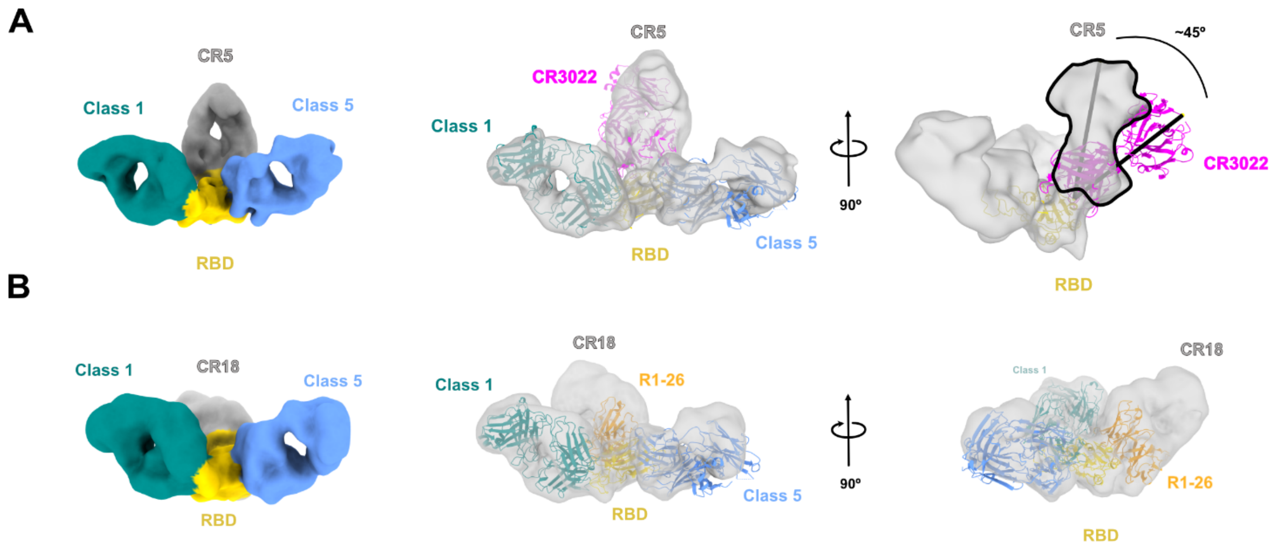

**Figure S7. Comparison of antibody angles of approach to the Class 4 cryptic RBD epitope.**
**(A)** Low-resolution reconstruction of the CR5 map with RBD in yellow, representative Class 1 in green, Class 5 in blue, and CR5 in dark gray. The CR3022 structure (PDB 6W41) is superimposed and shows an approximately 45° difference in angle of approach.
**(B)** Low-resolution reconstruction of the CR18 map with RBD in yellow, representative Class 1 in green, Class 5 in blue, and CR18 in light gray. The R1-26 variable domain (PDB 8ZHG; orange) is superimposed and shows a similar angle of approach and heavy- and light-chain orientation as CR18.

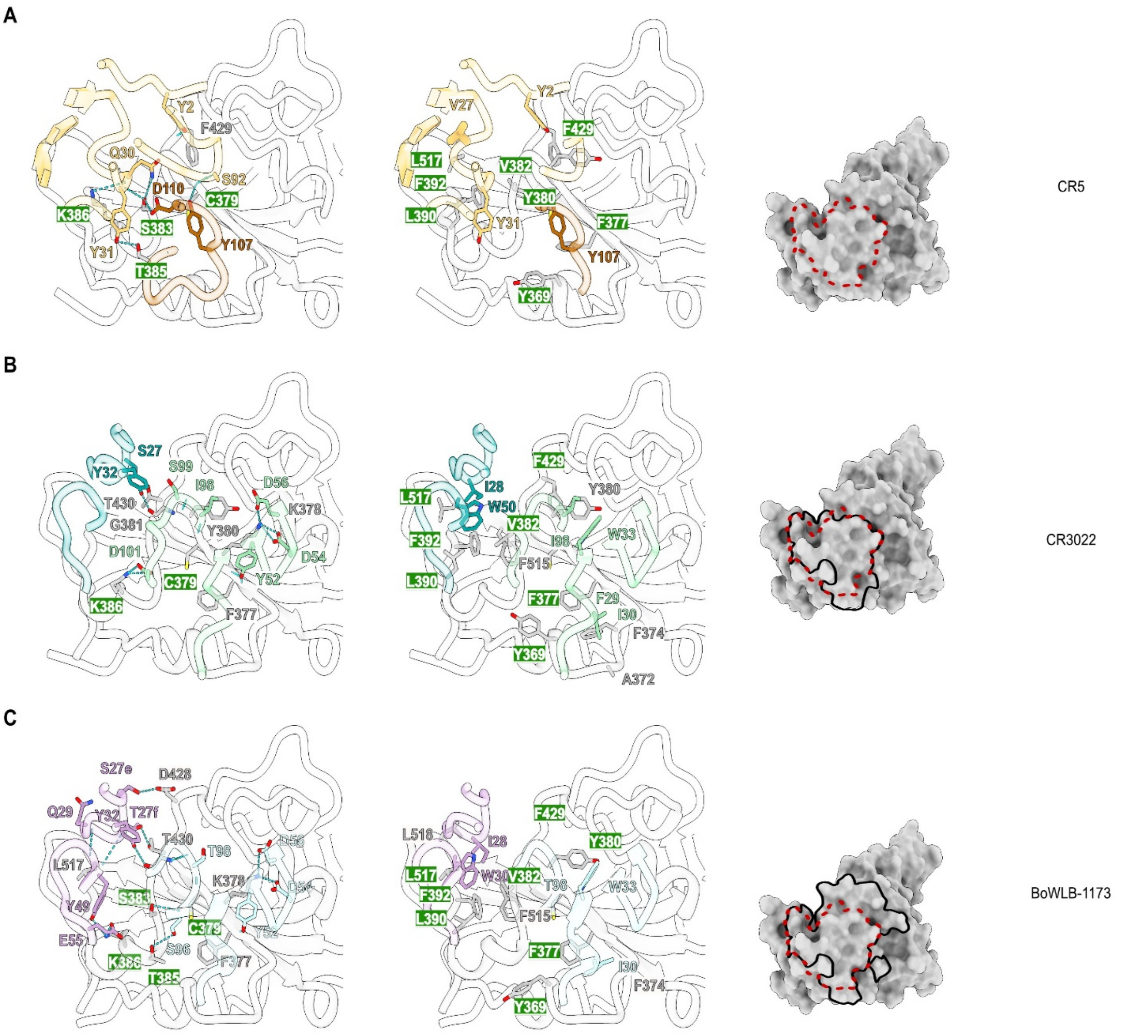

**Figure S8. Comparison of the binding modes of CR5, CR3022, and its affinity-matured CR3022 variant BoWLB-1173.**

**(A)** CR5 heavy and light chains are shown in yellow and maroon, respectively.

**(B)** CR3022 (PDB 6W41) heavy and light chains are shown in blue and green, respectively.

**(C)** BoWLB-1173 (PDB 9PSP) heavy and light chains are shown in blue and green, respectively. RBD is shown as a gray ribbon (left) or surface representation (right).

Epitope residues contacted by CR5, CR3022, and BoWLB-1173 are highlighted in green. The CR5 footprint is indicated by red dotted lines on the right.

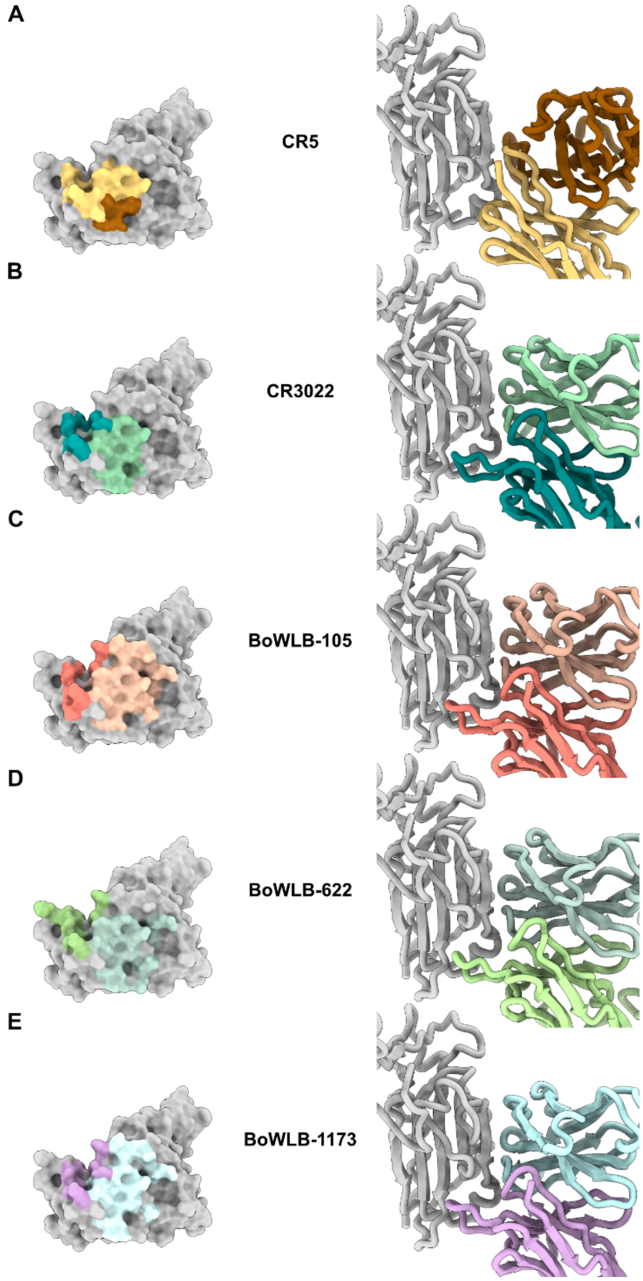

**Figure S9. Comparison of footprints and paratope organization of antibodies targeting the Class 4 cryptic epitope.**

**(A)** CR5, with light and heavy chains shown in yellow and maroon, respectively.
**(B)** CR3022, with light and heavy chains shown in pale green and blue, respectively.
**(C)** BoWLB-105, with light and heavy chains shown in pale pink and salmon, respectively.
**(D)** BoWLB-622, with light and heavy chains shown in pale green and chartreuse green, respectively.
**(E)** BoWLB-1173, with light and heavy chains shown in pale blue and purple, respectively.
For each antibody, the left column shows the epitope footprint on the RBD, while the right column compares paratope organization and angle of approach.

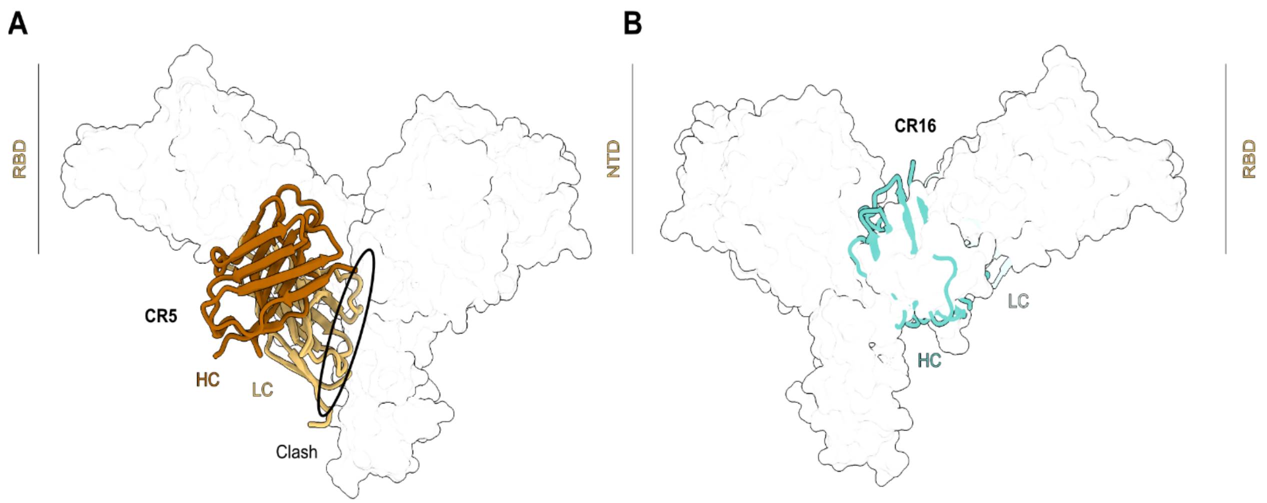

**Figure S10. Assessment of cryptic epitope accessibility on the soluble S1 subunit. (A)** CR5 and **(B)** CR16 models were aligned and superimposed onto the soluble S1 structure (PDB 7S0E). The CR5 light-chain variable domain partially overlaps with the NTD (highlighted in the black ellipse), whereas the heavy chain access the epitope without steric clashes. In contrast, the CR16 variable domain completely overlaps with the RBD.

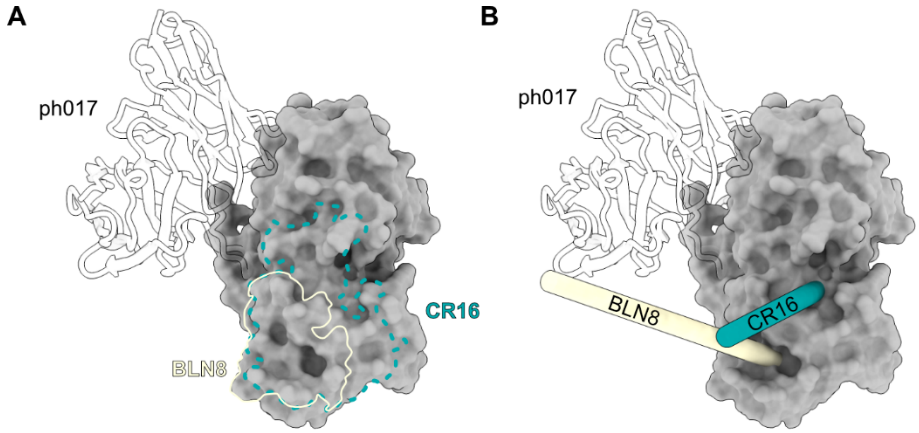

**Figure S11. Comparison of antibody footprints and angles of approach to the NTD cryptic epitope.**
**(A)** NTD surface is shown in gray. The CR16 footprint is outlined by a blue dotted line, and the NTD-8 footprint (PDB 9XAW) by a thin cream line.

**(B)** The principal axis of the NTD-8 (cream) and CR16 (blue) Fabs are represented as elongated cylinders to highlight their distinct angles of approach.

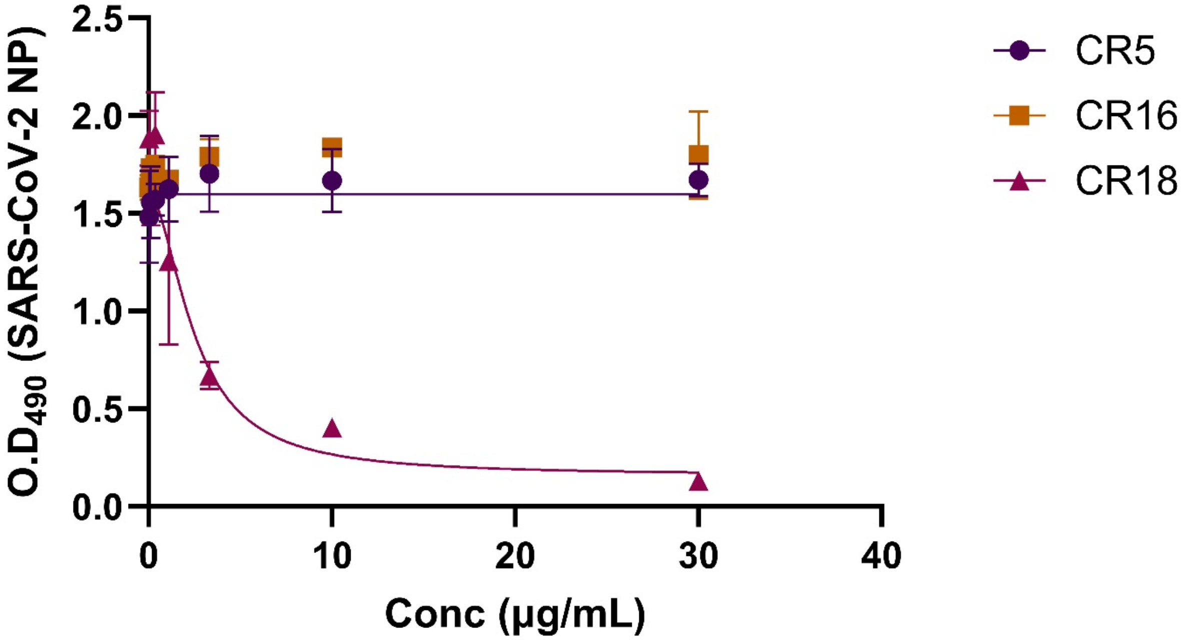

**Figure S12: Live-virus neutralization assay of CR5 (purple), CR16 (orange) and CR18 (pink) against WA1/2020 SARS-CoV-2.** Curves were fitted using the Absolute $IC_{50}$ function on GraphPad Prism.

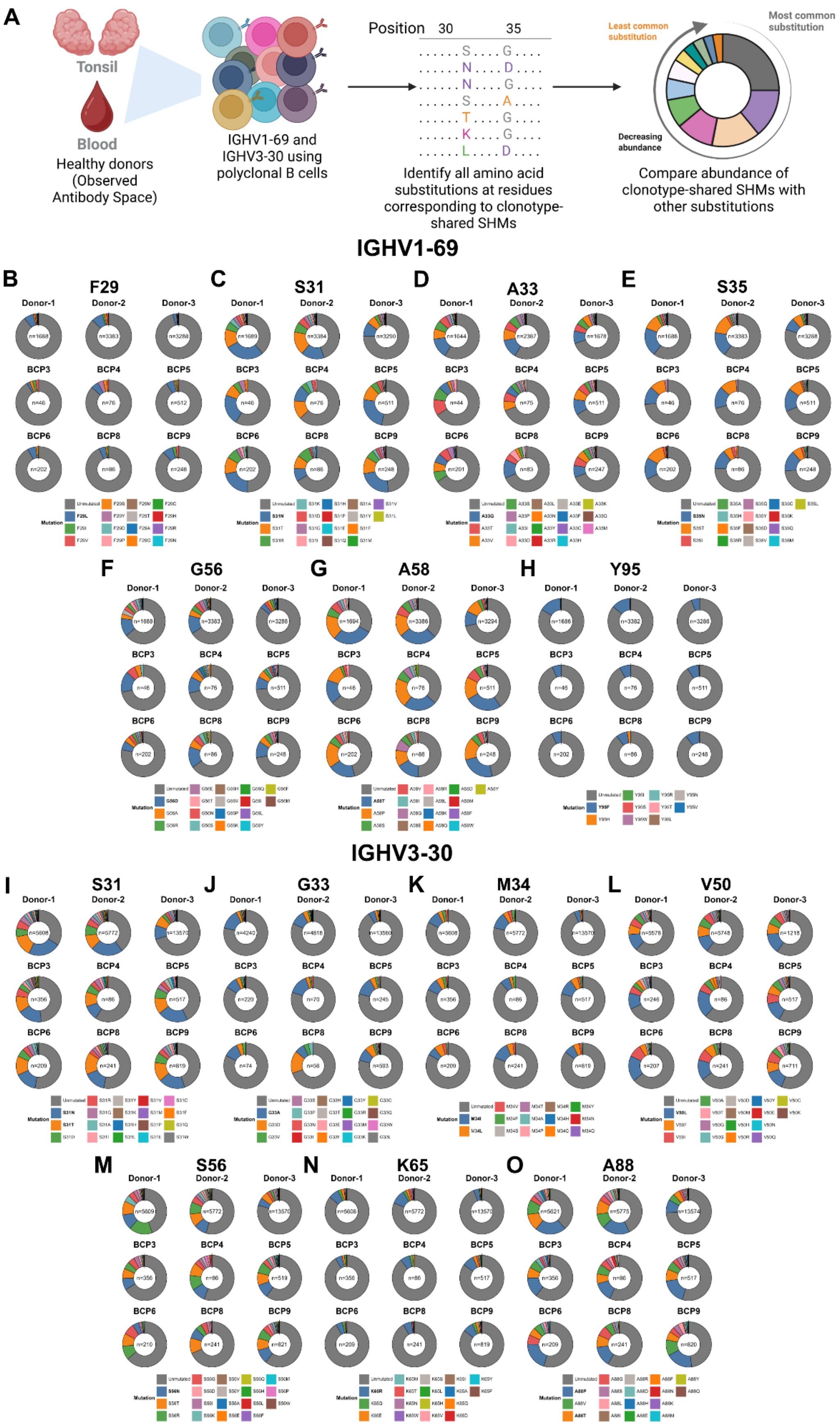
A
Tonsil
Blood
Healthy donors (Observed Antibody Space)
IGHV1-69 and IGHV3-30 using polyclonal B cells
Position 30 35
Identify all amino acid substitutions at residues corresponding to clonotype-shared SHMs
Least common substitution
Most common substitution
Decreasing abundance
Compare abundance of clonotype-shared SHMs with other substitutions
IGHV1-69
B F29
C S31
D A33
E S35
F G56
G A58
H Y95
IGHV3-30
I S31
J G33
K M34
L V50
M S56
N K65
O A88
Donor-1
Donor-2
Donor-3
BCP3
BCP4
BCP5
BCP6
BCP8
BCP9
Mutation
Unmutated

**Figure S13: Relative frequencies of different amino acid substitutions at sites with clonotype-shared SHMs.**

**(A)** Schematic representing the workflow used to define substitution frequencies.

**(B-H)** Donut plots depicting the relative frequencies of amino acid substitutions at F29, S31, A33, S35, G56, A58 and Y95 positions (left to right) in IGHV1-69-using BCR sequences

**(I-O)** Donut plots depicting the relative frequencies of amino acid substitutions at S31, G33, M34, V50, S56, K65 and A88 positions (left to right) in IGHV3-30-using BCR sequences.

Each segment represents an observed substitution and the size of the segment is proportional to frequency of the substitution in the respective individual. Substitutions are arranged clockwise in decreasing order of frequency. Substitutions representing clonotype-shared SHMs at each position are labelled in bold in the legend.

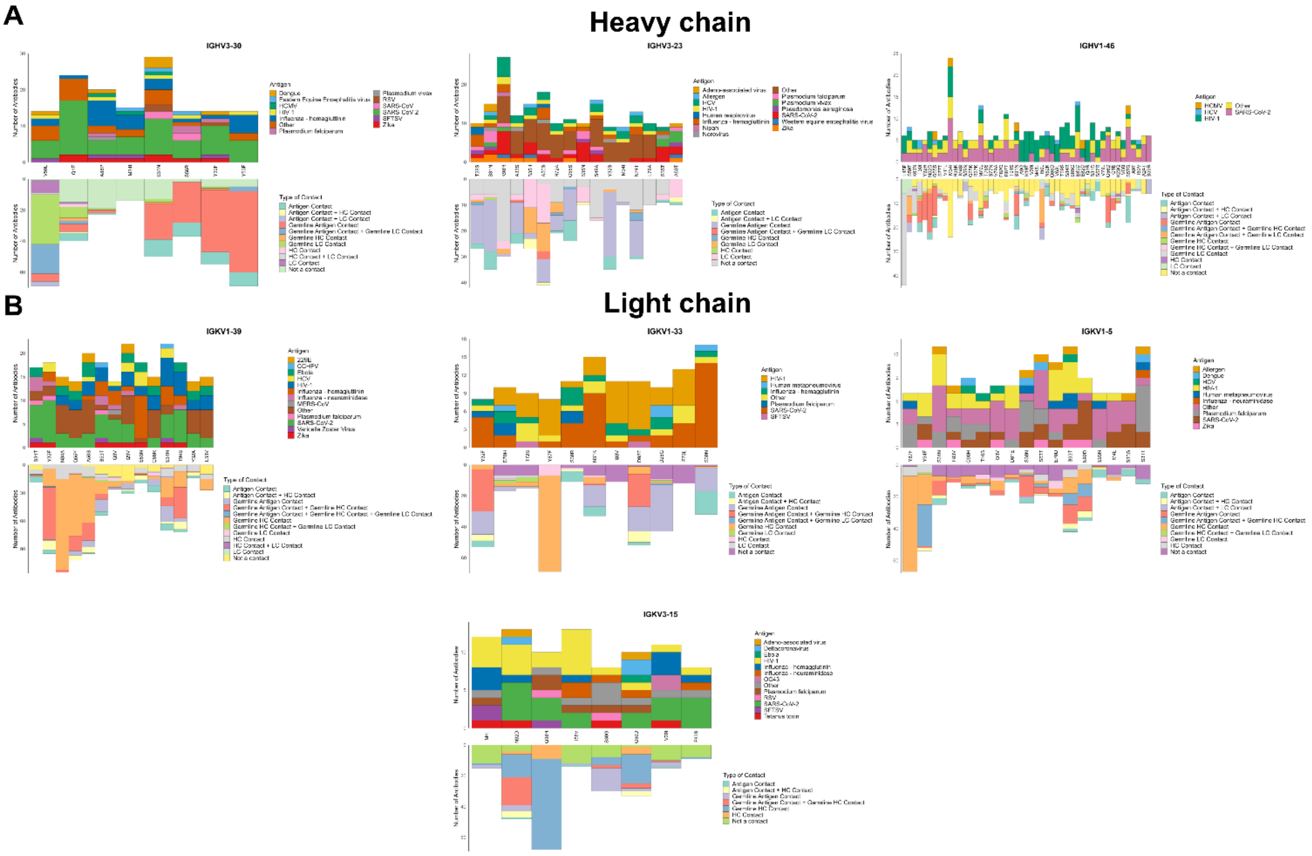

**Figure S14: Contributions of pan-antigenic SHMs to antigen–antibody interactions** Distribution of antigen specificities recognized by antibodies harboring a given pan-antigenic SHM (top panels), and the types of contacts formed either by the mutated residue or by the corresponding germline residue when the mutation is absent (bottom panels), for pan-antigenic SHMs in **(A)** heavy chain V genes IGHV3-30, IGHV3-23, IGHV1-46 (left to right) and **(B)** light chain V genes IGKV1-39, IGKV1-33, IGKV1-5, IGKV3-15 (left to right).

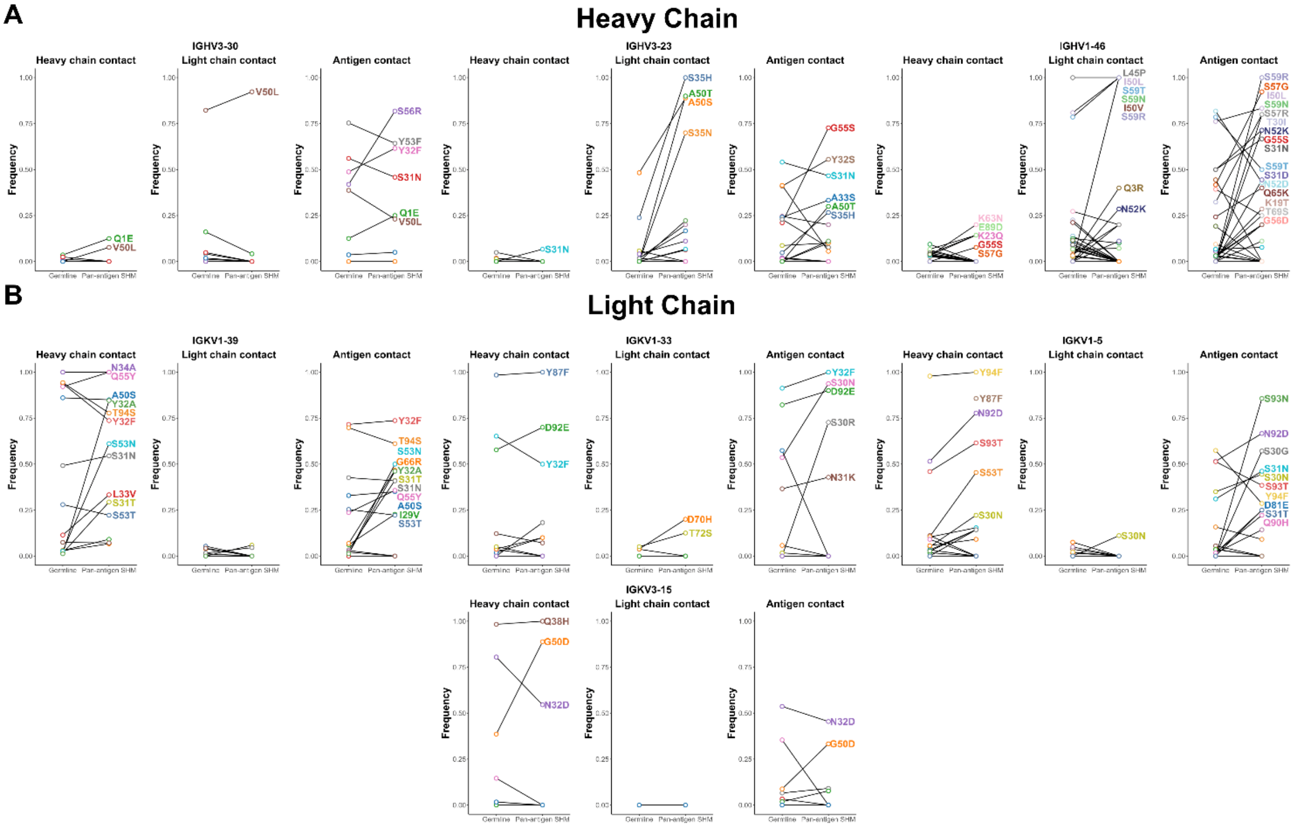

**Figure S15: Contact frequencies of pan-antigenic and germline contacts with cognate chain and antigen. (A-B)** Contact frequencies for pan-antigenic SHMs in **(A)** heavy chain V genes IGHV3-30, IGHV3-23, IGHV1-46 (left to right) and **(B)** light chain V genes IGKV1-39, IGKV1-33, IGKV1-5, IGKV3-15 (left to right). Contact frequency is defined as the proportion of antibodies in which a given residue, either mutated or germline, forms a contact with the heavy chain (left), light chain (center), or antigen (right). Linked dots represent paired comparisons of contact frequency for the same residue in its germline versus mutated (pan-antigenic SHM) state. Each color corresponds to a distinct pan-antigenic SHM.

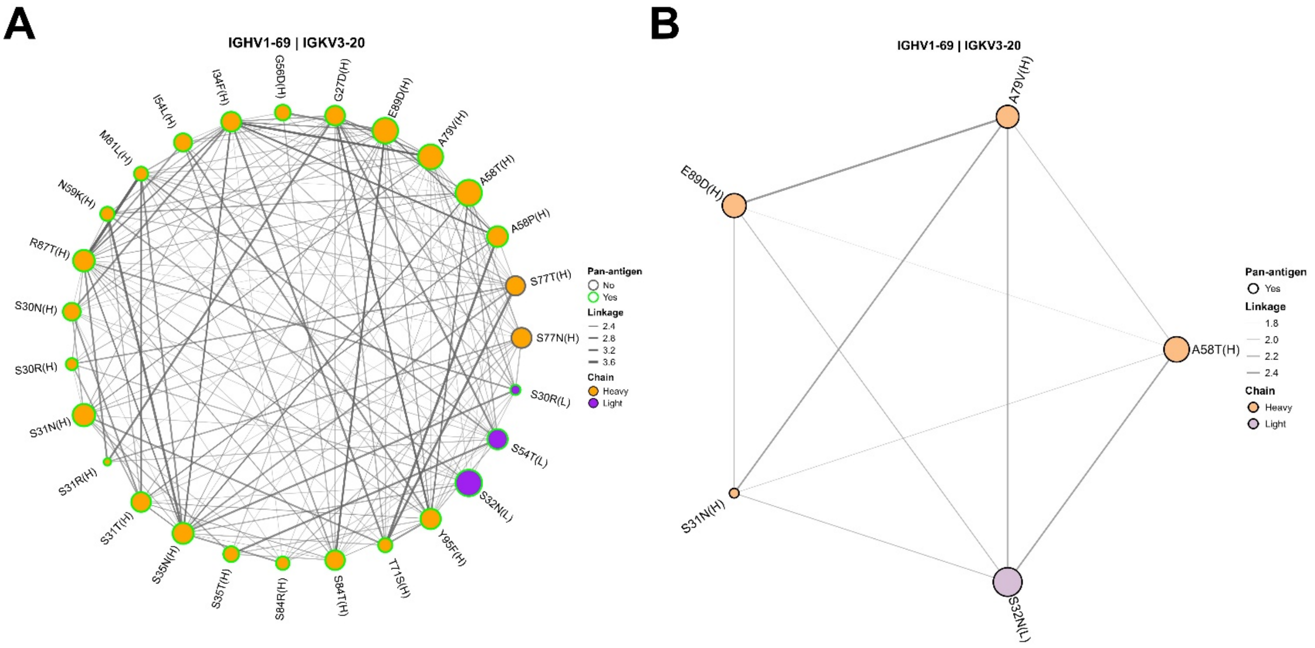

**Figure S16: Linkage analysis between pan-antigenic SHMs and other SHMs in IGHV1-69/IGKV3-20-using B cells obtained from healthy donors (OAS). (A and B)** Circos plot depicting linkage **(A)** between pan-antigenic SHMs and SHMs with linkage greater than 2 and **(B)** between pan-antigenic SHMs and SHMs observed in at least 10 percent of sequences, among non-naïve B cells with at least one SHM. Mutation nodes are colored based on the chain (orange – heavy, purple – light) and the outline node color indicates whether the mutation is a pan-antigenic SHM (green) or not (grey). The thickness of edges is proportional to the extent of linkage between the mutations.
